# Orphan genes shape the genome and parasitic arsenal of root-knot nematodes

**DOI:** 10.64898/2025.12.19.695360

**Authors:** Ercan Seçkin, Dominique Colinet, Marc Bailly-Bechet, Aurélie Seassau, Silvia Bottini, Edoardo Sarti, Etienne GJ Danchin

## Abstract

Orphan genes, lacking detectable homologs in other species, are common across eukaryotic genomes and can arise through divergence of existing genes or *de novo* from non-coding regions. Here we identified and characterized orphan genes in eight root-knot nematode species (genus *Meloidogyne*), the most destructive plant-parasitic nematodes. For that, we used comparative genomics across 85 nematodes, ancestral sequence reconstruction, synteny analyses, and multi-omics data. We found that around 16% of *Meloidogyne* genes are genus-specific transcribed orphan genes, with about 20% resulting from high divergence and 18% emerging *de novo*, often in transposon-rich genomic regions. Transcriptomic, translatomic, and proteomic evidence confirmed expression and translation of many orphan genes, which tend to encode shorter, secreted proteins. Notably, orphan genes are preferentially expressed during the infective juvenile stage and contribute substantially to the arsenal of nematode parasitism effectors. These findings highlight orphan genes as a significant and dynamic component of *Meloidogyne* genomes, potentially underpinning their parasitic adaptation and success.

## Introduction

### Orphan genes and *de novo* gene birth

In eukaryotes, genomes systematically contain a number of genes (up to 30%) that have no identifiable homolog in other species, known as orphan genes ^1,2^. The emergence of orphan genes is an important evolutionary mechanism that can be associated with the acquisition of functional novelty. Orphan genes may originate from pre-existing genes that have accumulated high divergence, reaching the point of no recognizable homology. This process can be facilitated by gene duplication or horizontal gene transfer events, followed by rapid evolution. Studies in several species suggest, however, that this explanation applies only to a subset of existing orphan genes ^3^. Another, non-mutually exclusive hypothesis is that orphan genes may emerge from non-genic regions. This phenomenon, known as *de novo* gene birth, occurs when previously non-coding and/or non-transcribed DNA sequences acquire the capacity to be transcribed and translated into a functional protein ^4–6^. For a long time, *de novo* emergence was considered highly unlikely because (i) it was believed that a new gene can only emerge from an existing gene and (ii) the probability that a newly emerged gene coding for a functional protein would be maintained in populations by selection was considered extremely low ^7^. However, with the expansion of genomic sequencing projects and the resulting increase in available genome data from a higher diversity of species, it has become clear that *de novo* gene emergence is not as rare as initially thought, and that many species- or lineage-specific sequences lack recognizable homologs ^8^. Several studies have taken advantage of this expanded set of genome data to confirm the existence of genes that likely emerged *de novo* ^1,9,10^.

In the case of a protein-coding gene, *de novo* emergence involves two main complementary processes: (i) transcription of initially non-coding DNA, and (ii) acquisition of an open reading frame (ORF). The relative order of these events allows two mechanisms to be distinguished ^10^: “transcription first” and “ORF first”. However, it is important to note that the distinction between the “transcription first” and “ORF first” mechanisms is not always straightforward. Just as it can be difficult to definitively classify an orphan gene as *de novo* or highly diverged, the temporal sequence of transcription and ORF acquisition may not be neatly separated. For instance, an ORF formed in a region of low transcription can gradually acquire regulatory features. Alternatively a *de novo* gene can later undergo rapid divergence, which can obscure its origin.

### Studies of orphan and *de novo* genes

Orphan genes have been identified in many species, and their proportion in the genome varies depending on the species ^2^. For instance, orphan genes represent 1.3% of genes in humans ^11^, 2.3% in *Drosophila melanogaster* ^12^, 4.4% in *Caenorhabditis elegans* ^13^, 25% in *Saccharomyces cerevisiae* ^14^, and 33% in *Pristionchus pacificus*, a nematode closely related to *C. elegans* ^15^. However, these numbers are barely comparable because studies have been made at different time points and by using different methodologies.

Concerning the origin of orphan genes, *de novo* emergence has been described in several species ^10^. This phenomenon was first reported in *D. melanogaster* in 2006 ^16^ and in *S. cerevisiae* in 2008 ^17^. Subsequently, it was estimated that 12% of orphan genes in *D. melanogaster* ^18^ and 5.5% of orphan genes in humans ^11^ have probably emerged *de novo*. However, these figures seem low in light of a more recent study that estimates most orphan genes emerged *de novo* and suggests that there are *de novo* genes yet to be identified in these species ^3^.

Little is known about the function of *de novo* genes. In yeast, one study showed that a *de novo* gene encodes a protein involved in the DNA repair pathway during the stationary phase of *S. cerevisiae.* This protein contributes to the yeast’s robustness when placed in a nutrient-poor environment ^17^. In plants, the first *de novo* gene whose function has been characterized is QQS, an *Arabidopsis thaliana* gene identified in 2009 that is involved in carbon and nitrogen metabolism ^19^. In codfish, a *de novo* gene encoding an antifreeze glycoprotein has been identified ^20^. Finally, a human-specific *de novo* gene encoding a protein found in the brain has been described and appears to be associated with Alzheimer’s disease ^21^. In a recent review, we observed that many orphan genes in animals, and more specifically those that emerged *de novo*, are involved in reproductive functions or are specifically expressed in gonads ^2^.

### Nematodes of the genus *Meloidogyne*

Nematodes are arguably the most numerous animals on Earth, paralleled only by arthropods, and are found in every biotope ^22^. In addition to their prevalence, they include free-living species as well as animal and plant parasites which cause substantial health problems and have important economic impact. So far, studies of orphan genes in nematodes have been restricted to the model species *C. elegans* and *P. pacificus*, which are both free-living species and relatively closely related. Orphan genes have not yet been investigated in parasitic nematodes, despite their importance for health and economy.

Overall, more than 27,000 nematode species have been described, and grouped into 12 main clades ^23^. Of these, four include plant parasites, ^24–26^, which collectively cause over $157 billion of economic loss every year to the global agricultural production ^27^. The most problematic of these worms regarding agriculture are the root-knot nematodes of the genus *Meloidogyne*, which belongs to Clade 12 (*Tylenchida*) ^24^. In addition to their economic importance, these species display peculiar biological features, such as diverse reproductive modes and ploidy levels ^28,29^. *Meloidogyne* species have been classified in three main clades (I-III), based on ribosomal DNA markers ^30^. Most species in Clade I are polyploid and reproduce exclusively via parthenogenesis, while in the other clades, most species are diploid and able to switch between parthenogenesis and sexual reproduction.

*Meloidogyne* species hijack plant root development to induce giant cells from which they feed, leading to gall formation on plant roots and causing billions of dollars in damage to agricultural production each year. Their life cycle, lasting three to six weeks, is divided into two phases: an exophytic phase outside the plant and an endophytic phase within the plant roots. The exophytic phase begins with the J1 juvenile larval form (within the egg), and is followed by the J2 stage, responsible for penetrating the root and initiating the endophytic phase. This phase includes the J3 and J4 larval stages, which are difficult to distinguish, followed by the transition of the larva to the adult stage. Adult males exit the root, while females remain sedentary, producing hundreds of eggs that are deposited on the root surface.

The study of *Meloidogyne* genomes began in 2008 with the sequencing of the *Meloidogyne incognita* genome ^31^. Early comparisons with the few available nematode genomes at the time suggested that a large fraction of *M. incognita* genes lacked detectable homologs in other species and were therefore considered species-specific. Subsequent improvements in genome assemblies and comparisons with additional nematode genomes confirmed the presence of thousands of such genes in *Meloidogyne*, although their evolutionary origin remained largely unexplored ^28,32^.

More recently, a comparison of gene sets from seven *Meloidogyne* species with those of 56 other nematodes, followed by a homology search against the NCBI’s non-redundant (nr) library, revealed tens of thousands of proteins specific to *Meloidogyne* species in *M. incognita* ^33^. Most of the corresponding genes were supported by transcriptomic data. While the origin of these numerous orphan genes was questioned, it was not investigated whether they highly diverged from ancestral sequences or emerged *de novo*.

Overall, these preliminary studies hypothesized that these numerous lineage-specific genes, supported by transcriptomic data, might play important roles, including for plant parasitism. Consistent with this view, these nematodes secrete proteins into host plants that act as effectors of parasitism, most of which are encoded by genes lacking identifiable homologs, i.e, orphan genes ^34^. Orphan effector genes involved in parasitic success have attracted a lot of attention as potential targets for more specific nematode control strategies. However, despite their importance, the origin, evolutionary history, and prevalence of orphan genes in *Meloidogyne* genomes remain unstudied.

Given their significant economic impact, unique biological features, and the previously reported abundance of candidate orphan genes, investigating the evolutionary origin, fate, and characteristics of orphan genes in these species, as well as their influence on genome structure and biology is timely.

In this study, we identified orphan genes in eight *Meloidogyne* species, including the most economically important ones (*M. arenaria, M. chitwoodi, M. enterolobii, M. graminicola, M. hapla, M. incognita, M. javanica,* and *M. luci)*. This dataset covers the three main *Meloidogyne* clades and spans an estimated ∼65 My of evolution since their last common ancestor ^35^. By comparing their sets of predicted proteins with those of 85 other nematodes and proteins in the NCBI’s nr database, we found that 16% of proteins lacked homology in other species to date and were encoded by transcriptionally-supported genes. Using ancestral sequence reconstruction and conserved synteny analysis with outgroup species, we inferred that at least 20% of the orphan genes most likely arose *de novo* from non-coding regions. In *M. incognita*, we found that DNA transposons were more prevalent around *de novo* genes compared to evolutionarily conserved ‘old’ genes. Furthermore, Meloidogyne-specific genes were more expressed at the second juvenile (J2) stage, which is responsible for plant recognition, penetration, and establishment of the feeding structure. Finally, orphan proteins tend to be shorter and more frequently secreted than the rest of proteins. Overall, these observations support a substantial contribution of orphan genes to the arsenal of effector genes involved in manipulation of plant defense.

## Materials & Methods

A diagram summarizing the entire workflow is shown in Figure 1.

**Figure 1:**
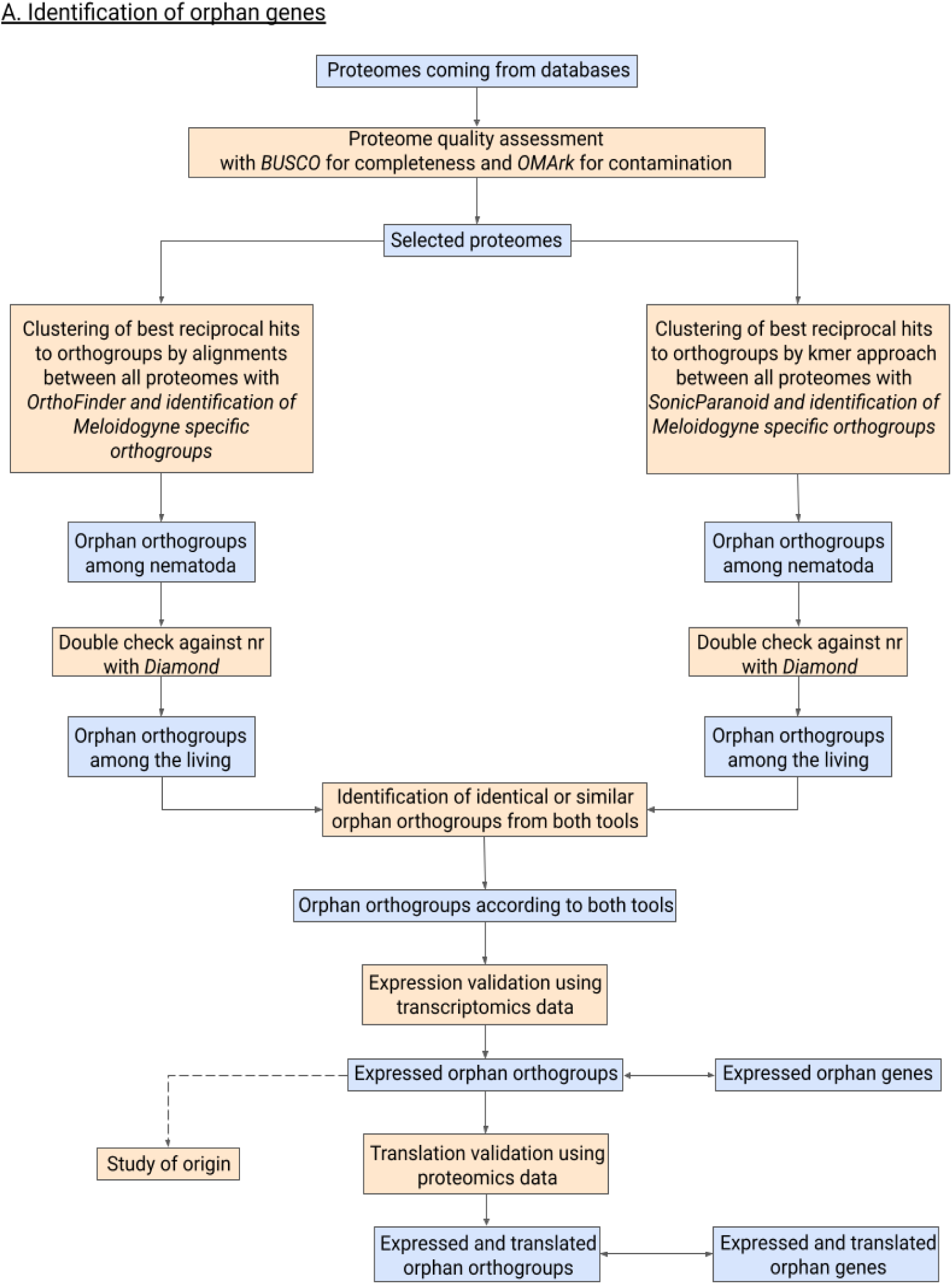
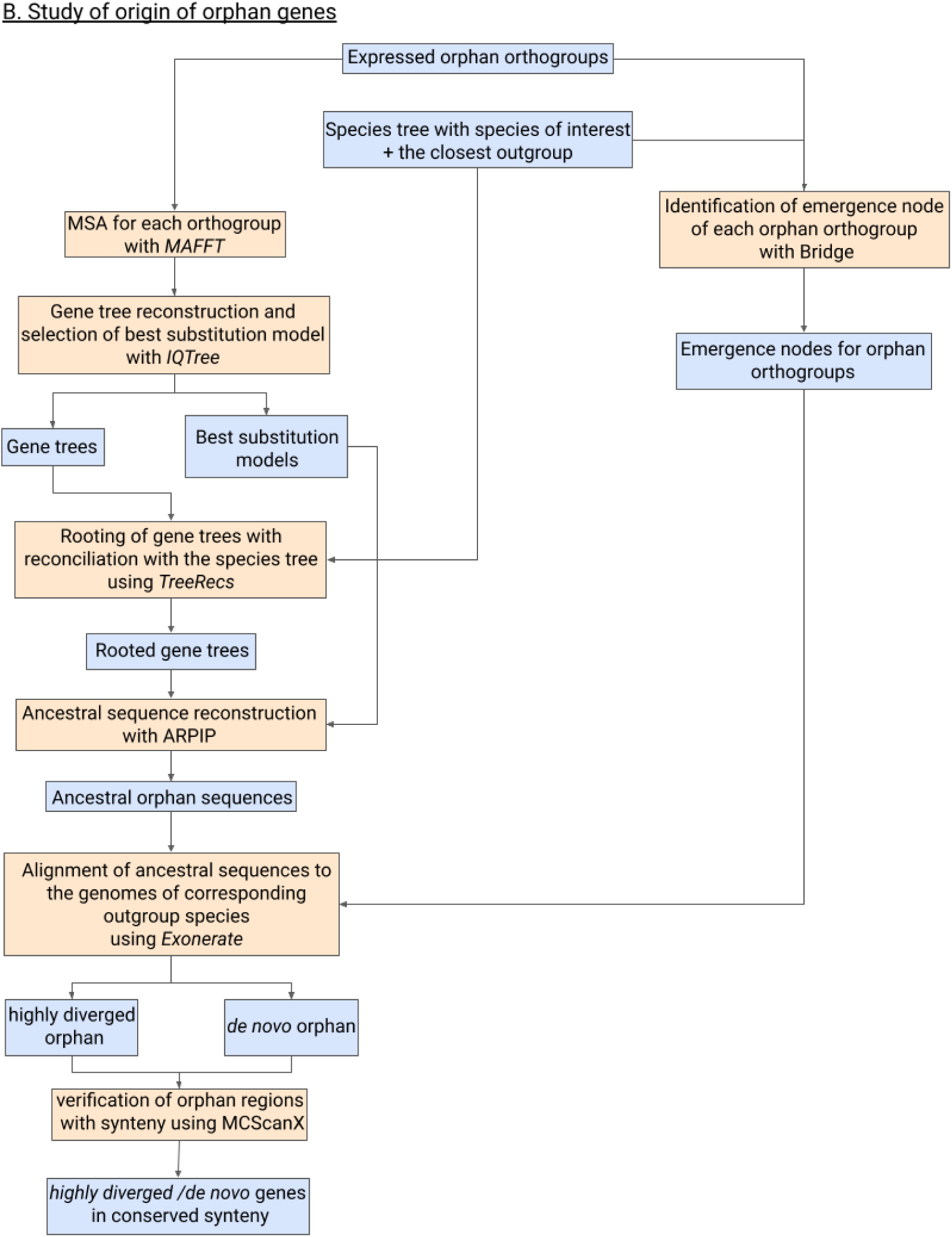
Bioinformatics pipeline to identify orphan genes and study their evolution in *Meloidogyne*. The different steps of the built pipeline are shown sequentially. Blue boxes correspond to inputs or outputs, whereas orange boxes correspond to different steps or analyses. A. First part of the built pipeline, used to identify orphan genes in *Meloidogyne*. B. Second part of the pipeline, used to study the origin of the identified orphan genes and to classify them as either highly diverged or *de novo*-emerged genes.

## 1. Selection of nematode predicted proteomes

A total of 206 nematode predicted proteomes are available in the WormBase Parasite database, including different *Meloidogyne* species ^36^. We made a selection from all proteomes based on two criteria: (i) quality of annotation and (ii) diversity of species represented. To measure the quality of the proteomes, completeness was assessed using BUSCO ^37^ (v 5.4.4, with metazoa_odb10 as dataset). This tool calculates the percentage of ‘universally’ conserved genes in a branch of the phylogenetic tree (here *metazoa*) within each proteome and yields a BUSCO score corresponding to the percentage of conserved genes actually found in the proteome. Absence of contamination in the selected proteomes was then assessed with OMArk ^38^ that calculates possible contaminations on the proteome by flagging a protein as contaminant if a protein has high similarity to sequences from non-target taxa. We defined a minimal BUSCO completeness threshold of 62% and a maximum OMArk contamination of 5%. Concerning the criteria of diversity of species represented, all species of the genus *Meloidogyne*, which is the group of interest, were retained if their proteome satisfied the BUSCO and OMArk criteria for completeness and lack of contamination. For the other genera, a maximum of three species for which the proteome met the above quality criteria were retained for each genus to provide a balanced representation in the dataset of selected proteomes.

## 2. Comparative analysis of predicted proteomes in Nematoda

Two different tools were used to perform the comparative analysis of the 85 nematode proteomes that were selected in the previous step. One of these tools is OrthoFinder ^39^ (v 2.5.4) that performs a homology search of all proteomes against all proteomes using Diamond ^40^, a tool similar to BLAST ^41^ but much faster due to its use of a simplified protein alphabet and binary data to align sequences. OrthoFinder was utilized with default parameters, using dendroblast ^42^ for gene tree inference and setting an e-value of 0.001. Based on this homology search between all provided protein sequences, OrthoFinder groups the best mutual hits into orthogroups. For each orthogroup, OrthoFinder generates a phylogenetic tree to identify orthologs. The proteome of the tardigrade *Hypsibius exemplaris* was used as a nematode outgroup to help root the OrthoFinder-reconstructed trees. The second tool used was SonicParanoid2 ^43^, which is analogous in its methodology but relies on a different clustering method, using graphs to identify homologs rather than MCL clustering and phylogenies, as OrthoFinder does. SonicParanoid uses MMSeqs in sensitive mode to align sequences all against all using k-mers ^44^. These two tools were chosen mainly to have a range of alignment strategies, one being based on BLAST-like heuristic method and the other utilizing k-mers.

OrthoFinder and SonicParanoid both produce two main output files. The first file is a matrix of protein counts per species for each orthogroup. The second file contains the orthogroup identifiers and the proteins that belong to these groups. Using a custom python script, orthogroups containing exclusively proteins from *Meloidogyne* species were selected for both tools. To reduce the risk of including proteins that would be false-predictions of annotation software, all orthogroups comprising a single *Meloidogyne* species (or a single predicted protein-coding orphan gene) were isolated from the orphan dataset. Therefore, the final set of retained orthogroups contained proteins present in at least two *Meloidogyne* species and lacking detectable homologs in the other species.

Following the homology search (see below), the orthogroups inferred by the two tools were compared. We classified orthogroups as identical, similar, and subset using a home-made script that compares the elements of each orthogroup between them. Identical orthogroups contained exactly the same set of proteins in both tools and were retained as such. Similar orthogroups shared at least 75% of their proteins with another orthogroup. For these, a union merged orthogroup was constructed. Finally, subset orthogroups contained only a portion of the proteins from an orthogroup identified by the other tool. In such cases, the largest orthogroup was retained. All retained proteins were considered encoded by potential *Meloidogyne*-specific orphan genes lacking detectable homologs in other nematodes.

## 3. Homology search among living organisms

A homology search was performed for the above-mentioned *Meloidogyne*-species orphan proteins using Diamond ^40^ against the NCBI non-redundant (nr) protein database. The e-value threshold was set to 0.001 and a minimum sequence identity of 25% was required. The more-sensitive option was used to enhance the sensitivity of the analysis and approximate BLASTp results. The top 100 hits for each query were used to infer a taxonomic annotation. Diamond was run in last common ancestor (LCA) mode to assign taxonomic labels to the sequences based on the NCBI taxonomic tree. The protein list for each *Meloidogyne* species was obtained, along with the taxon ID of the last common ancestor of their hits for each protein query. A homemade python script was used with the ete3 package ^45^ to convert taxon ID numbers into taxon names. Next, we eliminated orthogroups that contained at least one protein having a significant hit outside the genus *Meloidogyne*.

## 4. Transcriptional support for orphan genes

Short-read paired-end RNA-seq ^46^ transcriptomic data were available for *M. incognita* ^47^ (data for four developmental stages in triplicates: egg (W), second stage infective juvenile (J2), stage 3-4 juvenile (J3-J4) and adult female), *M. graminicola* ^48^ (data for four developmental stages in triplicates: W, J2, J3-J4 and adult female), *M. luci* (data for a mixture of stages with replicates in different conditions), *M. enterolobii* ^49^ (data for two developmental stages in triplicates: J2 and J3-J4), *M. arenaria* ^47^ (data for a mixture of stages without replicates), *M. javanica* ^47^ (data for a mixture of stages without replicates). These data were aligned to the reference genome of each corresponding *Meloidogyne* species using STAR ^50^. Then, RSEM ^51^ was used to determine expression values in FPKM (Fragments Per Kilobase Of Exon Per Million Fragments Mapped) for each gene. Thereafter, the results were normalized while maintaining positive expression values. The use of a base-10 logarithm allowed the data to be scaled without distorting the differences in value ranges or resulting in a loss of information. It is noteworthy that the addition of a value of 1 to the initial value yields positive expression values or zero, but never negative values.

When no replicates were available, the first quartile for expression values was calculated. Among the orphan genes in these species, those with expression values higher than this first quartile, calculated for the total gene set, were considered expressed. When replicates were available for specific stages or conditions, the median expression value for each gene was determined, also with the first quartile as a threshold to consider the gene as expressed. Finally, when a given gene was found to be expressed in at least one of the stages or conditions, it was considered to be expressed in the species under study.

For the analysis of developmental stages at different evolutionary ages, for *M. incognita* expression values, TPM results were used for each stage at each replicate. For each stage, median of the replicates were used and these medians were compared between different stages to assess the stage where the expression is at its maximum.

## 5. Ribo-seq support for orphan genes

In addition to the transcriptomic evidence, we incorporated translatomic ribosome profiling (Ribo-seq) data from a recent study ^52^ on *Meloidogyne incognita* to provide initial support for translation. The dataset included three developmental stages: pre-J2, J3–J4, and adult female, each with three biological replicates. Following the original authors’ approach, we processed cleaned Ribo-seq reads: by aligning the reads to the *M. incognita* genome using STAR ^50^ (--outFilterMismatchNmax 2, --outFilterMatchNmin 16, --alignEndsType EndToEnd) and performed transcript quantification with RSEM ^51^, yielding FPKM values. Then, we treated each FPKM table of each developmental stage of each replicate in the exact way that we treated transcriptomics data. We normalized the tables and the first quartile of each FPKM value for each replicate is used as criteria to consider a gene as translated or not. When a given gene was found to be translated in at least one stage or condition, it was considered to be translated. Finally, if there was at least one gene with evidence for translation in an orthogroup, all genes of the orthogroup were considered as translated.

## 6. Proteomic support for orphan proteins

To further support the existence of orphan proteins, we analyzed mass spectrometry-based proteomics dataset from the same study on *M. incognita* ^52^. The dataset consists of three different stages in three replicates: female, juvenile 2 and juvenile 3-4. We used the PEAKS Studio software ^53^ to identify predicted proteins with peptides matching with this mass spectrometry data for *M. incognita*. This analysis was applied not only on the predicted proteome of *M. incognita* but also on those of closely related Clade I species *M. luci, M. javanica* and *M. arenaria.* The other Clade I species*, M. enterolobii,* was excluded because it is too distant from the others and does not share the same sub-genomes ancestry.

We considered a protein as detected if at least one peptide matched significantly in any condition. To ensure reliability, we filtered results based on statistical significance, retaining only those with p-values below 5% and false discovery rate (FDR) below 1%.

We then identified orphan orthogroups containing at least one protein with proteomic support. As for transcriptomic and translatomic analyses, if a protein with mass spectrometry evidence belonged to an orphan orthogroup, we considered all proteins within that orthogroup to be supported by proteomic data.

## 7. Reconstructing the emergence dynamics of orphan genes

Two different tools, Bridge ^54^ and PastML ^55^, were used to estimate the ancestral presence/absence status of each orthogroup along each branch of the species phylogenetic tree, in order to investigate their emergence dynamics. PastML uses its own maximum likelihood (ML) method, called MPPA, which accounts for branch lengths in the species tree. Bridge also implements a dedicated algorithm, in which gene loss is more likely than gene gain. Both analyses were performed using the species tree inferred by OrthoFinder, based on homology relationships. To ensure the accuracy of our species tree, we compared it with a recently published mitochondrial tree for *Meloidogyne* species ^48^ and confirmed that the topologies were congruent. Bridge, implemented in R, was run with input data formatted as a table containing three columns (protein ID, species name, and orthogroup ID), along with a rooted species tree. The output consisted of a table listing orthogroups, nodes, and associated p-values. PastML was executed using its command line version, with input data structured as a matrix with orthogroups as columns and species as rows, along with a rooted species tree including branch lengths. The output was a tabular file containing node IDs and the presence of orthogroups at those nodes. The results from each tool were then filtered using a python script to identify the node of first occurrence of each orphan orthogroup along the branches of the species tree. Finally, the inferred number of emerging orthogroups at each node of the species tree was compared between the two methods.

Our dataset includes four species in Clade I that share recent common ancestry and parental A and B subgenomes (*M. incognita*, *M. luci*, *M. javanica* and *M. arenaria*). Our initial inclusion criterion for orphan genes was their presence in at least two *Meloidogyne* species. Therefore, retention is more likely in this group than the rest of *Meloidogyne* species. To assess whether this could artificially inflate the number of emergence events along the branch leading to this group, we performed two additional analyses. In the first analysis, we kept only *M. incognita* as a representative of this group. In the second analysis, we kept only *M. incognita* and *M. luci*, the two most closely related species. The emergence node of each orphan orthogroup was then recalculated by Bridge under these two conditions.

For the analysis of developmental stages at different evolutionary ages, Bridge was used to identify the emergence node of transcriptionally validated orthogroups of *M. incognita,* using the same parameters as before, but with *M. incognita* as the unique reference species. We classified orthogroups into three evolutionary age classes: orphan, intermediary, and old. Old orthogroups correspond to genes that are inferred to have emerged at the two deepest nodes, whereas intermediary orthogroups include all nodes between these and the *Meloidogyne-*specific orphan genes. The reference species tree was obtained from the OrthoFinder tree, with modifications to the positions of higher-level taxonomic groups. Although OrthoFinder correctly resolves relationships among species within clades, the relative positions of some families and orders were inconsistent with previously established phylogenies ^24^. Therefore, after removing branch lengths (as Bridge does not use them), we repositioned these higher-level groups to match the reference topology.

## 8. Identification of potential *de novo* genes specific to the *Meloidogyne* genus

### 8.1 Ancestral sequence reconstruction for each *Meloidogyne*-specific orthogroup

For each *Meloidogyne*-specific orthogroup, protein sequences were aligned using MAFFT ^56^. A maximum likelihood phylogenetic tree was then constructed for each orthogroup using IQ-TREE ^57^, using the mset option to evaluate and select the best-fitting evolutionary model among WAG, JTT and LG. These three models were chosen because they are the only ones available in both IQ-TREE and ARPIP (see below). Each tree was then rooted using TreeRecs ^58^ by reconciling it with the species tree. TreeRecs maps each gene in the gene tree to its corresponding species in the species tree and, by comparing the topology of the gene tree with that of the species tree, infers evolutionary events such as gene duplications and losses. TreeRecs then identifies the optimal root position in the gene tree that minimizes the number of inferred duplications and losses, ensuring a rooting consistent with the species tree. Finally, ancestral sequences for each orthogroup were reconstructed with ARPIP ^59^, using the best-fitting model identified by IQ-TREE. ARPIP was selected over alternatives such as RaxML ^60^ and IQ-TREE because it treats gaps as separate character states, allowing meaningful inference of insertion and deletion events, whereas other tools treat gaps as missing data.

### 8.2 Alignment of ancestral protein sequences on the outgroup species

For orthogroups that contain at least one gene from Clade III *Meloidogyne* species, the reconstructed ancestral protein sequences were aligned to the genome of *Pratylenchus penetrans*, the closest outgroup species with an annotated genome available, using Exonerate ^61^ (v 2.4.0). For other orthogroups shared by only a subset or clade of *Meloidogyne* species, the ancestral proteins were aligned to the genome of the closest outgroup *Meloidogyne* species using the same approach. Exonerate, using the protein2genome model, aligns protein sequences to a genome translated in the six reading frames while accounting for possible introns. We used the default minimum score threshold of 100 and retained the three best alignments for each ancestral protein sequence.

To investigate the mechanisms underlying the transition from a non-coding region to a transcribed coding gene, a home-made script was used to identify regions that aligned imperfectly according to Exonerate results. Specifically, we searched for premature stop codons, indels not in multiples of three, and incorrect splice sites in the outgroup genome, as these features are expected to disrupt correct transcription and translation.

Some genes detected in *Meloidogyne* may also exist in the outgroup species but remain unidentified due to potential gene prediction errors. To account for this, all ancestral sequences that aligned perfectly with the outgroup genome without any of the above problems were classified as potentially unpredicted genes in the outgroup. Using GFF3 annotation files for all genomes, we verified whether these alignments corresponded to the position of a predicted gene in the outgroup genome. Using this approach, orphan genes were classified as highly diverged if a gene was already predicted at the same locus but had no detectable homologs in the species harboring the orphan gene, or as *de novo* if no gene was predicted at the matching locus and the alignment exhibited disruptive features described above such as premature stop codons.

### 8.3 Synteny analysis

We assessed whether orphan genes within the *Meloidogyne* genus are located in conserved synteny contexts using MCScanX (48). This step was essential to identify regions that are highly diverged or that may have given rise to *de novo* genes. MCScanX identifies collinear syntenic regions by combining protein sequence similarity using BLASTp results with corresponding gene position via genome annotation files. MCScanX requires two concatenated input files, one containing all pairwise BLASTp results and another listing gene positions obtained from genome annotation files. These files were generated from BLASTp results and GFF3 annotations corresponding to different comparisons and across all *Meloidogyne* species and the closest outgroup species, *P. penetrans*. Since most *Meloidogyne* genomes in Clade I are triploid or tetraploid, MCScanX was run with default parameters for simplicity, but with -b inter-species to identify collinear blocks only between different species, an e-value of 0.001, and - s 4 to require at least four genes per collinear block. The tool produced a set of syntenic blocks along with a detailed list of collinear gene pairs.

Following the initial analysis, the pairwise collinearity output file was reformatted to focus on syntenic blocks containing orphan genes. By default, MCScanX only reports collinear gene pairs, meaning genes without collinear homologs are absent from the output. For example, if gene n and gene n+2 are collinear with y and y+1, respectively, the missing gene n+1 is absent from the output file. To address this, we developed a home-made python script to identify genes missing from syntenic blocks by pinpointing orphan genes on the collinearity file, enabling us to study the genes that flank orphans. This analysis provided additional support for orphan gene classification by mapping both collinear gene positions and missing genes in the syntenic blocks.

Finally, we compared the list of candidate highly diverged orphan genes and the list of *de novo* gene candidates with synteny results using a second home-made script. This allowed us to determine whether the flanking genes of the highly diverged or *de novo* gene candidates are conserved in synteny between the species harboring the gene and their outgroup.

### 8.4 Transposable elements

Transposons were detected using EarlGrey, with a 100 bp minimal size filter. The categories “Satellite”, “Unknown”, “Simple_repeat”, and “Low_complexity were discarded. Transposable elements (TE) annotated by Earlgrey were clustered according to their DFAM annotation before analysis; hence, all Tc1-Mariner elements, all hAT elements, and all Gipsy elements were grouped together. Transposons detected as “SINE?” were counted as “SINE”. Qualitative and statistical analysis were then performed in R.

## 9. Contribution of orphan genes to the parasitism

To assess whether orphan genes contribute to the set of parasitism effectors of *Meloidogyne* species, we retrieved a list of known effectors (a total of 111 effectors) from a previous study ^63^, in which the identified effectors meet the criterion that the corresponding genes are expressed in at least one secretion organ. We then used cdhit2D (Li and Godzik 2006), with the parameters -s2 0 -c 0.90 -g1 -aL 0.30 -aS 0.30, to cluster known effector protein sequences with the predicted proteomes from our study. Then, by cross-referencing the list of effectors with the cd-hit clusters we defined which and how many orphan proteins correspond to effectors.

To assess whether feeding tube proteins ^64^ are orphans, as this study was based on an older version of *M. incognita* genome, we retrieved the protein sequences of feeding tubes and performed blastp searches with default parameters against the proteome predicted from the updated version of the genome. Once the corresponding sequences in the current predicted proteome were identified, we used our orphan classification to determine whether these proteins should be considered orphans.

## 10. Characteristic features of orphan genes

Various properties of *Meloidogyne* orphan and non-orphan sequences were measured at both the gene and protein levels to enable comparisons and identify features characteristic of orphans.

At the protein level, we measured:

i. 22 physicochemical properties using a python script and the *Bio.SeqUtils.ProtParam* module from Biopython (27). These included the instability index, hydrophobicity profile, average flexibility, average hydropathy, isoelectric point, protein charge at pH 7.5, proportions of aliphatic amino acids and charged amino acids, molar extinction coefficient, cumulative proportions of small, aliphatic, aromatic, non-polar, polar, charged, basic and acidic amino acids in a helix, sheet or loop, and sequence length and molecular weight.
ii. The relative abundance of each amino acid.
iii. Three functional features: the presence of InterPro protein domains detected with InterProScan ^66^, the presence of secretory signal peptides detected with SignalP ^67^ (v 6.0), and predicted subcellular localization with DeepLoc2 ^68^.

At the gene level, we analyzed nucleotide frequencies and sequence length using a python script.

A summary table was created for each sequence by retrieving data from the output files of each script and tool (Supplementary Table 1). All these features were then compared between orphan and non-orphan sequences at both the protein and gene levels. Statistical tests were applied depending on the feature: Student’s t-test or Mann-Whitney test for quantitative features (according to normality calculated with Shapiro test), and Fisher’s exact test for qualitative features. P-values were reported for each feature, with values lower than 0.05 considered significant. Given the large dataset, even small differences might be significant. Therefore, for any feature that was significantly different, we calculated both relative and absolute differences in mean values between orphan and non-orphan sequences to assess whether the difference can be considered different enough to have a biological relevance.

Furthermore, to characterize the level of divergence among orphan and non-orphan proteins within orthogroups, pairwise percent identities between *Meloidogyne* proteins were calculated using Diamond. Orphan orthogroups selected for pairwise comparisons were those identified as expressed. Non-orphan orthogroups selected for pairwise comparisons were the ones for which all *Meloidogyne* species were present as well as at least five additional nematode species. Pairwise comparisons were then performed between species within orthogroups via a home-made script as followed: (i) among proteins from Clade I species, (ii) between proteins from Clade II species and those from the other clades, (iii) between proteins from Clade III species and those from the other clades. For non-orphan orthogroups, additional comparisons were performed between proteins from all *Meloidogyne* clades and those from *P. penetrans*. These percent ids were also taken into account to adjust parameters for ancestral sequence alignments on outgroup species.

Finally, rates of non-synonymous (Ka) and synonymous (Ks) mutations between orphan and non-orphan genes have been calculated using KaKs_Calculator ^69^ to compare their mutational retention patterns. As input, the tool requires CDS and protein sequences as well as pairs of homologs. As output, the software returns a file with Ka and Ks values between pairs of homologs. We calculated these values for pairs of homologs of orphans and non-orphans between *M. hapla* and *M. enterolobii*, *M. chitwoodi* and *M. hapla* to assess the levels of divergence across transitions from Clade II to Clade I and from Clade III to Clade II, respectively.

## 11. Classification of orphan genes and their origin using Random Forest

We trained two Random Forest classifiers ^70^ using the scikit-learn library (version 1.3.0) to assess whether orphan gene classifications could be predicted from their molecular characteristic features. We selected a subset of easily computable features (See data availability) from the full feature table. One classifier was built to distinguish between orphan and non-orphan sequences (as defined above), and the second to distinguish between highly diverged and *de novo* orphan sequences. For each task, we split the dataset into training and test sets, and performed hyperparameter optimization using GridSearchCV with 5-fold cross-validation. Model performance was evaluated on the held-out test set by standard classification metrics precision, recall, and F1-score. We extracted feature importances according to Gini impurity, using the built-in feature_importances_ attribute of the trained classifiers, to identify the variables that contributed most to classification. We also used shap tree explainer to determine the positive and negative impact of the features.

## Results

### 16% of *Meloidogyne* proteins are encoded by transcriptionally supported orphan genes

Nearly 1/4 of the predicted proteins lack homologs in other nematodes and their corresponding genes are supported by RNA-seq data

First, we performed a comparative analysis of predicted proteins across nematode species since most nematode species’ genomes are not represented in general public databases. From the available proteomes, we selected 85 representative nematode species using a quality criterion based on a minimum metazoa BUSCO score of 62% and the absence of contaminants according to OMArk, as well as a species representativeness criterion, covering a total of 44 genera and 10 clades (Supplementary Table 2). Of the nine *Meloidogyne* species for which a proteome was available, eight were retained, spanning the three main *Meloidogyne* clades (I, II, and III). Following an all-against-all sequence comparison, the best reciprocal homologs were identified and classified into orthogroups by OrthoFinder and SonicParanoid. OrthoFinder classified 92% of the 1,970,290 proteins from the 85 nematodes into a total of 275,056 orthogroups, while SonicParanoid classified 79% of all these proteins into 75,469 orthogroups. We then selected orthogroups that contained exclusively *Meloidogyne* proteins and excluded those that were specific to a single species in order to minimize the risk of species-specific gene prediction errors. This approach led to the identification of 15,024 and 14,955 orphan orthogroups according to OrthoFinder and SonicParanoid, respectively. These orthogroups respectively comprise a total of 81,914 and 64,342 orphan proteins, corresponding to 26% and 21% of the proteins predicted in *Meloidogyne* genomes. These *Meloidogyne*-specific proteins are therefore possibly encoded by orphan genes lacking detectable homologs in nematodes outside the genus *Meloidogyne*.

Although the presence of homologs in at least two *Meloidogyne* species minimizes the risk of prediction errors, it remains possible that the same error has been made by prediction software in multiple species. To further minimize such prediction errors, we retained only orphan proteins encoded by genes with substantial expression support from RNA-seq transcriptome data.

All *Meloidogyne*-specific orthogroups were analyzed accordingly, and only those containing at least one gene supported by transcriptomic data were retained. This analysis identified a total of 13,279 and 13,185 orthogroups (for OrthoFinder and SonicParanoid, respectively) that include at least one orphan protein whose corresponding gene is supported by transcriptomic data. This represents a total of 76,930 and 59,859 orphan proteins supported by expression data, respectively, encompassing 25% and 19% of all proteins predicted for the genus *Meloidogyne*.

### Most *Meloidogyne* orphan proteins relative to other nematodes also lack homologs in the rest of living organisms

The next verification step was to determine whether *Meloidogyne*-specific proteins remained orphans when considering all other living organisms beyond nematodes. We performed a homology search against the NCBI nr protein database, which spans all domains of life, to rule out two cases: (i) genes possibly acquired by horizontal transfer events between *Meloidogyne* and species other than nematodes, and (ii) genes inherited from an older common ancestor but lost in all nematodes except *Meloidogyne,* although this scenario seems less likely. Most proteins predicted to be orphans in *Meloidogyne* relative to other nematodes either had no hits with any proteins in the nr database or had hits only with proteins from *Meloidogyne* species. This result is not surprising, considering that nr includes proteomes from only a few *Meloidogyne* species. Only 273 proteins were found to have a significant hit outside the *Meloidogyne* genus and were therefore classified as non-orphans for further analysis. We then excluded all orthogroups identified by OrthoFinder and SonicParanoid that contained at least one of these non-orphan proteins, resulting in a total of 76,717 and 59,488 orphan proteins, corresponding to 24% and 19% of all predicted *Meloidogyne* proteins, respectively.

To ensure consistency in using both OrthoFinder and SonicParanoid to identify orphan proteins, we compared the results from the two tools after verifying transcription and confirming the absence of homology with other organisms. We compared all orphan orthogroups identified by both tools and retained those that either were identical, shared at least 75% of proteins, or were subsets of each other based on sequence composition. This approach left us with 8,974 non-redundant and transcriptionally verified orphan orthogroups supported by both methods. These comprised a total of 48,681 orphan proteins, which represent 16% of the *Meloidogyne* pan-proteome (Figure 2). Depending on the species, this proportion ranges between 6% to 21%, with the higher proportions observed in the parthenogenetic polyploid species from Clade I.

**Figure 2:**
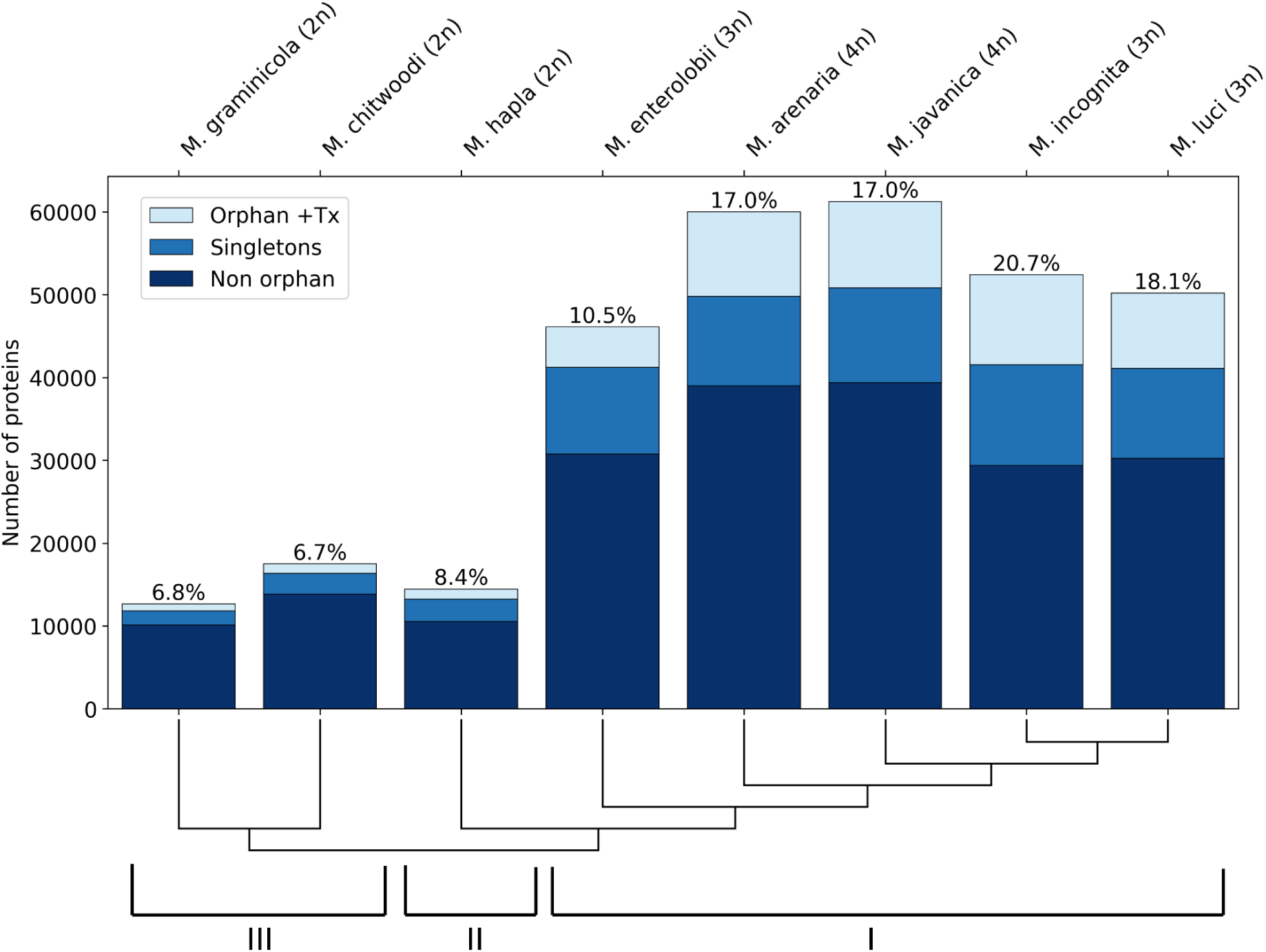
Number of orphan genes for the eight *Meloidogyne* species studied. Bar plot showing the number of orphan genes for each species. “+Tx” indicates transcriptomic support. The percentages above the bars correspond to the proportion of orphan proteins with transcriptional evidence for each species. A species tree illustrating clade relationships is shown below the plot.

### More than 30,000 orphan proteins are further supported by translatomics and/or proteomics data

Once transcriptionally-supported *Meloidogyne* orphan genes had been identified, our next objective was to assess evidence for translation. To do so, we used ribo-seq and proteomics data from a recent study on *M. incognita* ^52^. The ribo-seq data provided evidence for translation for 31,595 orphan genes. In addition, a total of 17,596 proteins from Clade I *Meloidogyne,* out of the 200,000 predicted proteins, were supported by proteomic data, regardless of their orphan status. After filtering for the orphans and considering that proteomic support within an orphan orthogroup indicates that the entire orthogroup is supported, we identified 1,134 orphan proteins whose existence is supported by proteomic data. When we took the union of supported translation either from ribo-seq or proteomics, we have in total 31,607 orphan proteins with evidence for translation. A venn diagram summarizing transcriptional and translational evidence is provided in the Supplementary Data (Supplementary Figure 1).

### The emergence of orphan genes in *Meloidogyne* shows several phases of acceleration

To understand the dynamics of orphan gene emergence in *Meloidogyne*, we estimated the most likely time of appearance for each transcriptionally-supported orphan orthogroup on the species tree, while accounting for potential subsequent gene losses in some species, using two different methods, Bridge and PastML (Figure 3).

**Figure 3:**
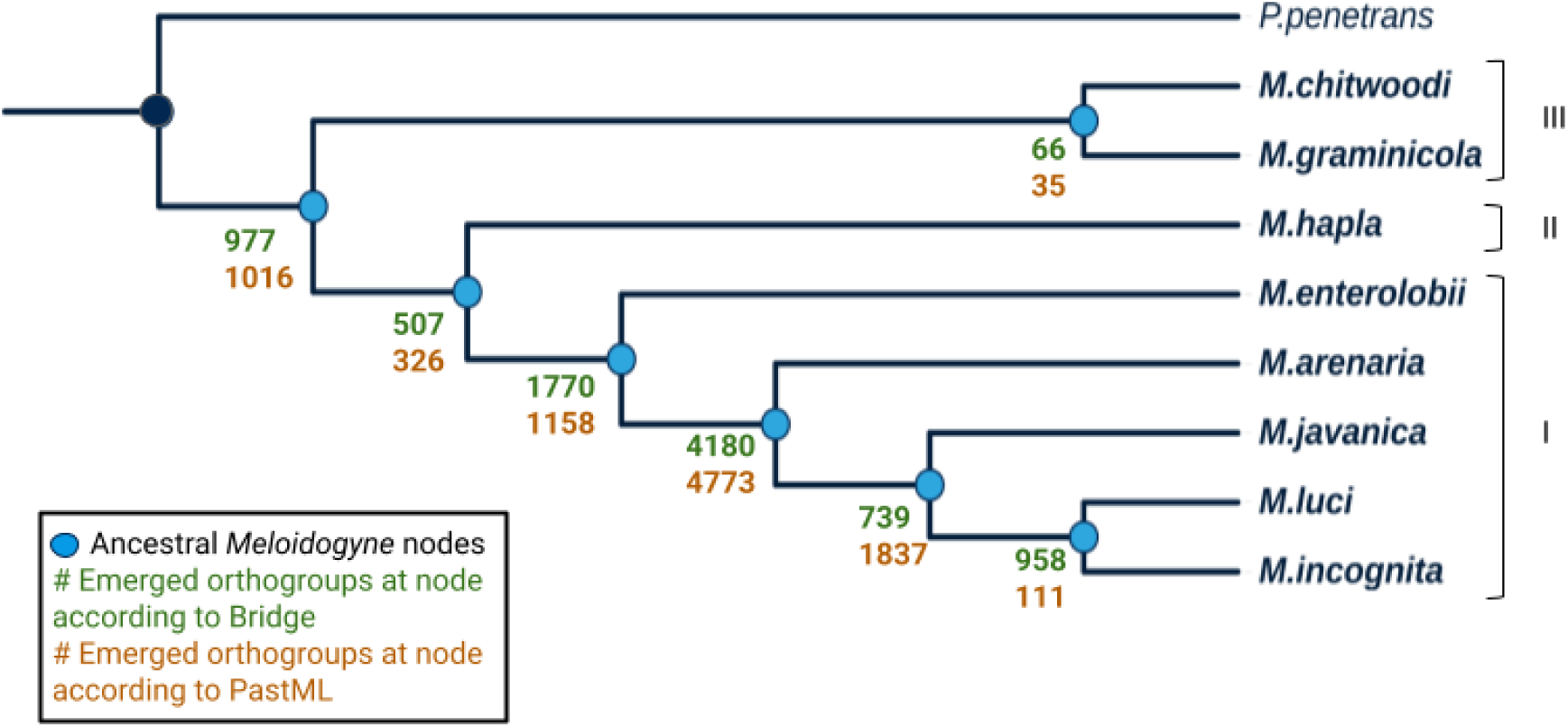
Number of emerged orphan orthogroups at each node of the *Meloidogyne* species tree, with *P. penetrans* as the closest outgroup. The species tree includes eight *Meloidogyne* species belonging to three different clades (indicated on the right) together with *P. penetrans*, the closest outgroup to this genus. Each ancestral *Meloidogyne* node is represented by a blue dot. The number of orphan orthogroups that emerged at each node is indicated next to it, in green for Bridge and in orange for PastML.

Despite some differences in the number of orthogroups predicted to have emerged at each node—particularly within Clade I (*M. javanica, M. luci,* and *M. incognita*)—the overall trend was highly similar between both methods. Our analysis revealed a steady increase in the number of orphan orthogroup emergence from the common ancestor to individual species. However, this increase was not uniform, as we identified several bursts of emergence, notably in the ancestor of Clade I and, more prominently, in the common ancestor of *M. arenaria, M. javanica, M. luci,* and *M. incognita*, where both methods predicted the highest number of emerged orphan, orthogroups (Figure 3).

It should be noted that these three species share common ancestral A and B subgenomes that they inherited from recent hybridization events ^47,48,71^. Therefore, if an orphan gene emerged ancestrally in A or B, it will likely be inherited by all the descendent species. Since one criterion for orphan gene retention in our analysis was presence in at least two different species, this has more chance to happen in this group of closely related species. *M. enterolobii*, on the other hand, has not been described as sharing the same A and B subgenomes, and according to mitochondrial-based phylogeny is quite distant from the rest of Clade I species ^47^.

This pattern aligns with observations from the species tree inferred by OrthoFinder (Supplementary Figure 2), where *M. enterolobii* appears more distantly related to other Clade I species based on branch lengths. Additionally, when analyzing the percent identities of orphan proteins within orthogroups, we found that orphan proteins from Clade I species (excluding *M. enterolobii*) shared a mean identity of 91.2% when compared among each other. However, when compared to *M. enterolobii*, this percentage dropped to 72%, confirming that *M. enterolobii* is substantially more divergent than the rest of Clade I species.

As presence of four closely related Clade I species might artificially inflate the number of orphan gene emergence compared to Clade II and III, we repeated the same analyses keeping only two or only one representative from these four species (Supplementary Figure 3). In both cases, the ancestral state reconstruction consistently confirmed a burst of orphan gene emergence in the branch leading to Clade I. These findings support the hypothesis of a burst of more than 1000 orphan gene families emerging in an ancestor of Clade I.

### Almost 20% of orphan genes in *Meloidogyne* emerged *de novo*, preferentially in genomic regions with higher density of DNA transposons

To identify the origin of orphan genes and distinguish *de novo* gene birth from high divergence from a pre-existing gene, comparison with an outgroup species and synteny analysis are essential. As a first step, we splice-aligned orphan proteins to the six-frame translation of the outgroup genomes lacking detectable homologs. To increase the likelihood of obtaining such splice alignments, we reconstructed ancestral protein sequences for each orphan orthogroup. In principle, these reconstructed ancestral sequences are phylogenetically closer to the outgroup. The *Pratylenchus* genus is the closest to *Meloidogyne,* and *P. penetrans* is the closest species with an available genome sequence ^24^. Therefore, for each orphan orthogroup that was deemed present in the last common ancestor of the *Meloidogyne* species (Figure 3), we splice-aligned the reconstructed ancestral protein to the six-frame translation of the genome of *P. penetrans*. For orphan groups that were not deemed present at the root, but instead specific to a *Meloidogyne* clade or branch, we splice-aligned the reconstructed protein to the six-frame translation of the genome of the closest outgroup *Meloidogyne* species.

Using gene annotation files for all genomes, we assessed whether inferred ancestral orphan proteins aligned to translation of genomic regions in outgroups that were compatible with the presence of a canonical gene. By definition, beyond the closest outgroup, orphan genes are absent from all other nematodes and even the rest of species. First, we observed that around 10% of ancestral reconstructed sequences aligned almost perfectly to the translation of the outgroup genome, in a manner compatible with proper transcription and translation. These were therefore considered false positive orphans, likely corresponding to genes that have not been predicted in the outgroup species. Using conserved synteny analysis based on flanking genes, we found that for around 20% of orphan genes, a different gene was present at the expected locus in the outgroup. This suggests that these genes may have strongly diverged from a common ancestral gene, impairing identification of homologs.

To identify genes that may have emerged *de novo* among the 70% remaining orphans, we examined genomic regions in the outgroups that matched with the ancestral sequence but displaying features incompatible with gene splicing or translation, such as premature STOP codons, frameshift-inducing indels (not in multiples of three), or incorrect splice signals. Mutations in such non-genic regions may have enabled the progressive emergence of coding sequences. We identified at least 2,031 ancestral orphan proteins that aligned to translation of outgroup genomes but in a manner incompatible with a functional gene (see example in Figure 4). The corresponding 8,833 genes likely emerged *de novo* in *Meloidogyne* species at different evolutionary time points, corresponding to 18% of orphan genes in this genus. These results suggest that *de novo* gene birth contributed substantially to the presence of orphan genes in the *Meloidogyne* genus.

**Figure 4:**
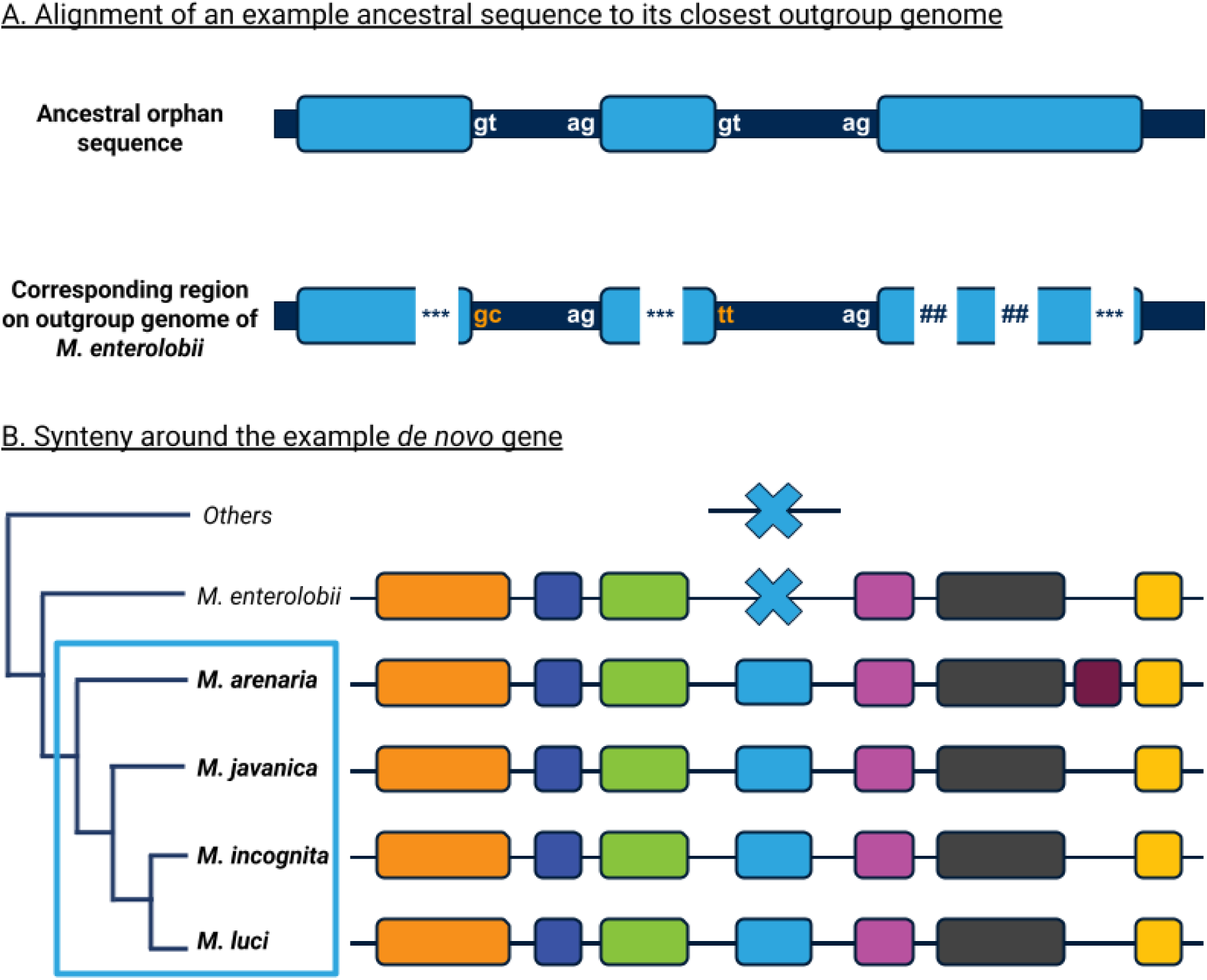
Example of a *de novo* gene birth event in Clade I. A. Alignment of the reconstructed ancestral sequence of an orthogroup common to all members of Clade I, except *M. enterolobii,* to the six-frame translation of *M.enterolobii* genome using Exonerate. Dark blue represents introns and light blue represents exons. STOP codons are represented by ***. Frameshift insertions or deletions are represented by #. Correct splicing sites are shown in white, and mutated ones in orange. B. Assessment of synteny conservation around the identified *de novo* gene. At the locus where the ancestral sequence aligns to the six-frame translation of *M. enterolobii* genome, the three closest upstream and downstream genes were examined to determine the conservation of the region. Lines represent intergenic regions, and boxes correspond to genes. Light blue boxes correspond to the *de novo* gene, and a blue cross indicates the corresponding locus in *M.enterolobii*. The color code indicates homologous genes across species. The blue rectangle on the species tree represents the species that contain the *de novo* gene.

The remaining ancestral orphan sequences could not be aligned to the closest outgroup translated genome. Therefore, for these cases, it is not possible to determine whether they correspond to highly diverged genes or *de novo* genes (Figure 5).

**Figure 5:**
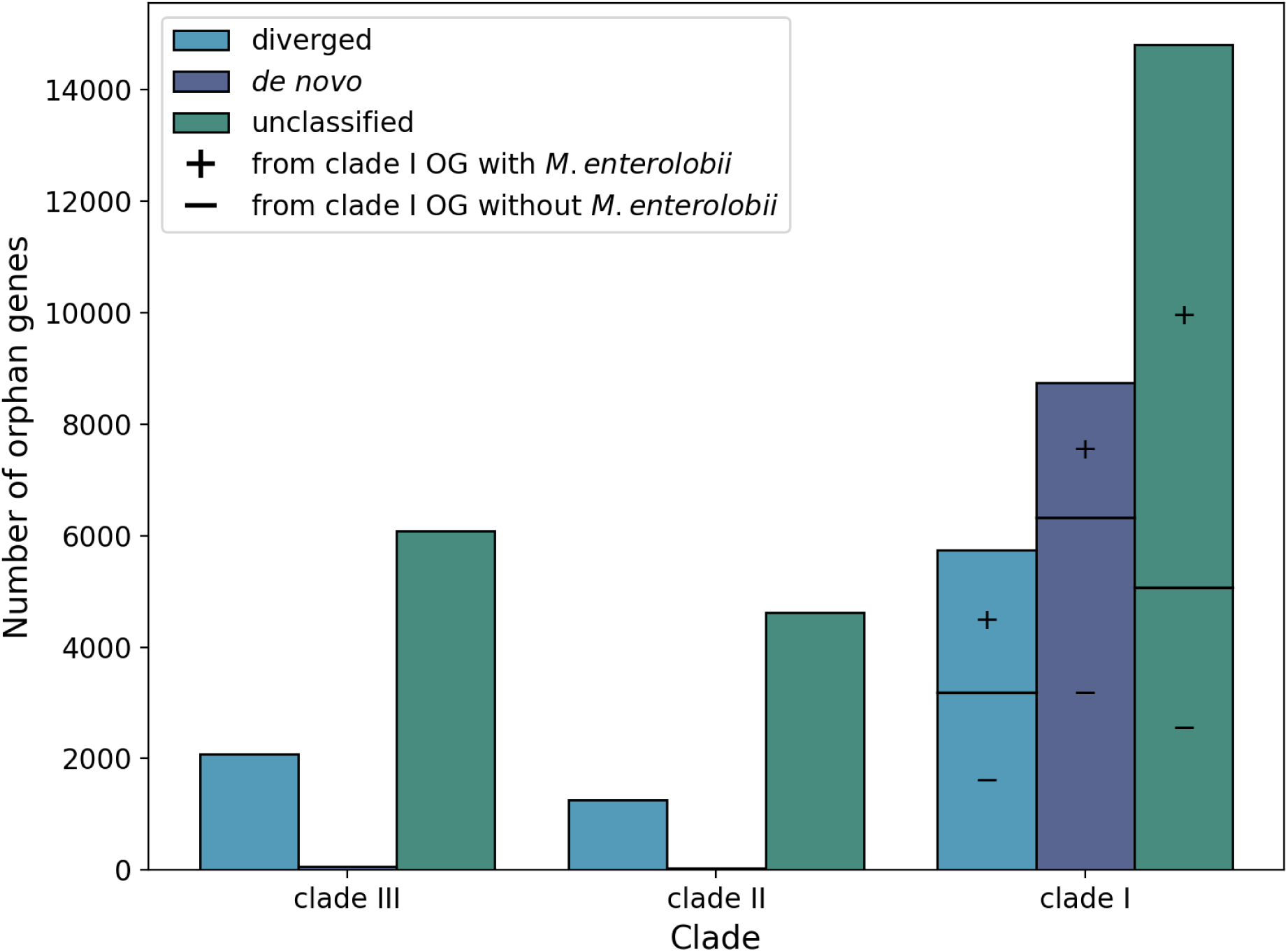
Number of orphan genes classified as highly diverged, *de novo,* or unclassified. Bar plots show the number of orphan genes for each clade and category. For Clade I species, orthogroups (OGs) are further subdivided based on whether they include *M. enterolobii* or not.

We further describe here an example of a likely *de novo* gene birth event involving an orphan gene common to all Clade I *Meloidogyne* species sharing A-B subgenomes, with the exception of *M. enterolobii.* In this case, the inferred ancestral orphan protein was aligned to the six-frame translation of the *M. enterolobii* genome, the closest outgroup, where the corresponding locus displayed multiple features incompatible with gene translation. Mutations that arose in an ancestor of the other Clade I species carrying the A-B subgenomes likely enabled the emergence of a sequence compatible with transcription and translation (Figure 4). As expected, the gene is also absent from *M. hapla* (Clade II), as well as from *M. graminicola* and *M. chitwoodi* (Clade III).

As transposable elements (TE) can trigger *de novo* transcription of genomic regions ^72^, we analyzed the co-distribution of TEs and *de novo* genes in the *M. incognita* genome. For each gene, we defined flanking regions as 1 kb upstream and 1 kb downstream, and we assessed the presence of at least one TE. We compared *de novo* genes with “old” evolutionarily conserved genes, defined as genes inferred to have emerged at the most ancestral nodes of the nematode tree of life. We observed a significant enrichment of TEs in the flanking regions of *de novo* genes (Chi-square test, *p* = 1 × 10⁻¹²). Specifically, 351 out of 2,258 *de novo* genes (15.5%) were flanked by at least one TE, compared to 2085 out of 19,767 old genes (10.5%). Among genes flanked by TEs, *de novo* genes were significantly more likely to be associated with DNA transposons (Chi-square test, *p* = 9 × 10⁻3), with 68.9% of TE-associated *de novo* genes flanked by a DNA TE, compared to 61.4% for old genes. At a finer taxonomic resolution, certain TE families were particularly enriched around *de novo* genes (Chi-square test, *p* = 8 × 10⁻^6^). Notably, DNA/PIF-Harbinger (4.0% vs. 1.6%) showed the strongest positive residuals when comparing *de novo* to old genes (Supplementary Figure 4). In contrast, SINE elements were strongly depleted around *de novo* genes, with only 0.05% of *de novo* genes flanked by SINEs compared to 5.1% of old genes. These patterns were consistent when analyzing upstream and downstream regions separately. Overall, these results suggest that *de novo* genes preferentially arise in TE-rich genomic regions, particularly in regions enriched in DNA transposons.

### Orphan genes contribute to the parasitic arsenal of root-knot nematodes and are preferentially expressed at the pre-parasitic stage

By comparing *Meloidogyne* predicted proteins to a reference dataset of 111 known parasitism effectors from a previous study ^63^, we recovered 83 clearly matching effectors under strict clustering parameters. Among these, 34 were encoded by orphan genes (including 9 singletons) representing approximately 40% of all matched effectors. This suggests that nearly half of the known parasitism-related effectors in *Meloidogyne* are of orphan origin. This finding is consistent with previous reports indicating that the majority of effectors secreted by nematodes are encoded by orphan genes ^34^.

A recent study identified the protein families involved in the formation of feeding tubes in root-knot nematodes ^64^. Feeding tubes are structures specific to root-knot nematodes and are crucial for proper feeding of the nematode from plant cell nutrients. We therefore investigated whether orphan proteins contribute to the composition of these structures. Our analysis showed that nearly all of these proteins are specific to root-knot nematodes, with only one protein family also detected in *P. penetrans*. These results further support a major contribution of orphan proteins to the specialized parasitism of *Meloidogyne* species.

Furthermore, we investigated how orphan genes are expressed during the life cycle of *M. incognita* in comparison with the other genes. We integrated stage-specific transcriptomic data from *M. incognita* with estimates of orthogroup age and inferred the emergence node of each transcribed orthogroup on the nematode species tree. This tree corresponds to a corrected version of the Orthofinder species tree, modified to fully match the reference topology of the nematode tree of life ^24^. We then classified the orthogroups into three evolutionary age classes: orphan, intermediate, and old. Old orthogroups correspond to genes inferred to have emerged at the two deepest nodes, whereas intermediate orthogroups include all emergence nodes between these and the *Meloidogyne-*specific orphan genes. For each gene belonging to these orthogroups, we then identified the developmental stage at which its expression was highest. The results showed that expression levels of genes belonging to old orthogroups are relatively similar at each stage and that the order of expression follows the temporality of development. In contrast, the expression of more recently emerged genes is increasingly biased toward the J2 stage at the expense of others (Figure 6). This indicates that recent genes are more likely to be preferentially expressed at the pre-parasitic J2 stage. This stage is crucial for the establishment of plant parasitism, as it is responsible for locating the plant roots, navigating to the roots, penetrating them and, once inside, establishing the feeding structure.

**Figure 6:**
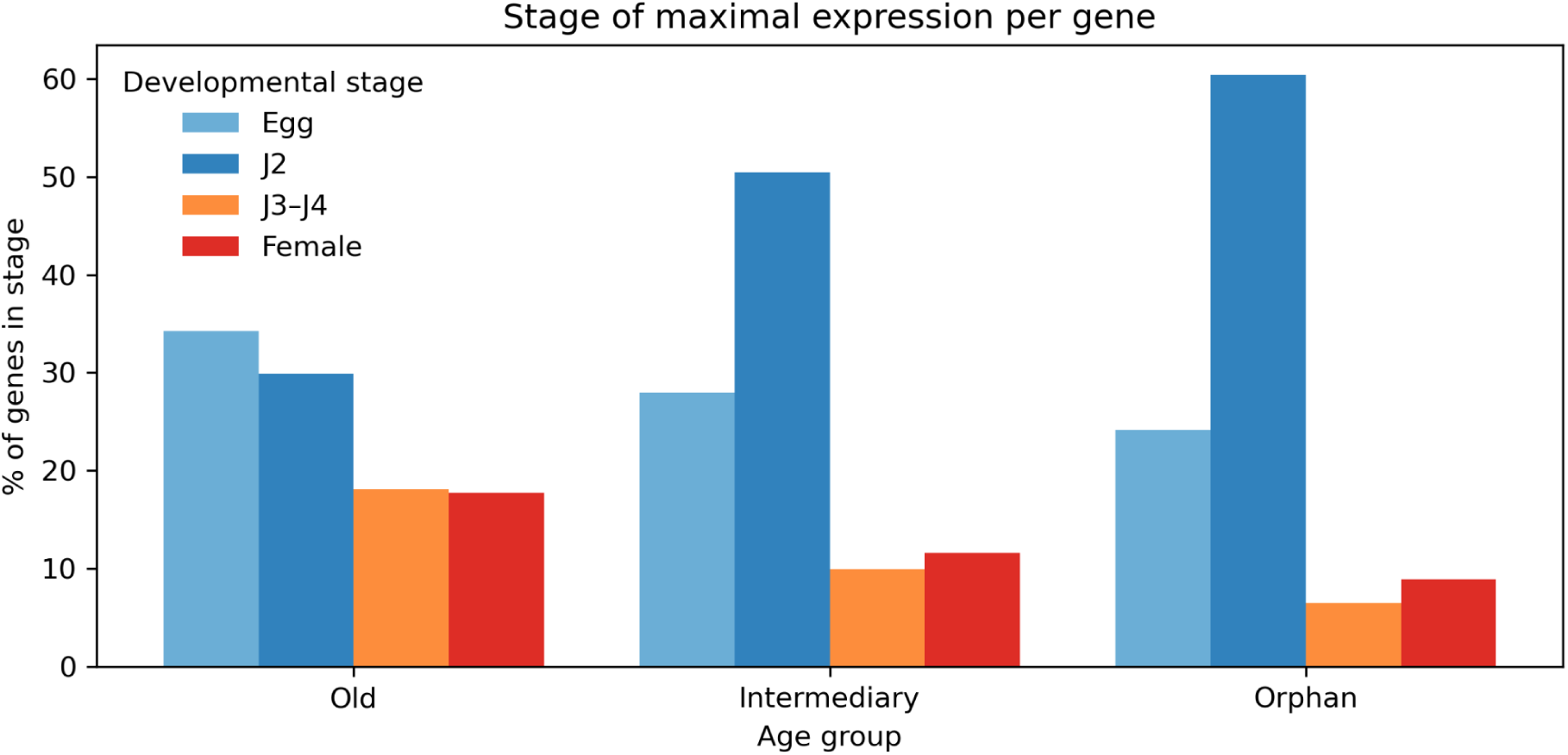
Developmental stage of maximal expression per gene. Genes were grouped according to their evolutionary age, based on orthogroup emergence along the species phylogeny. The old age group comprises genes inferred to have emerged at the ancestral node separating nematodes from their closest outgroup, *H. exemplaris*. Intermediate genes correspond to orthogroups that emerged at internal nodes between this ancestral split and the orphan group, with orphan genes representing genes specific to *Meloidogyne* genus.

### Orphan genes tend to encode shorter and more frequently secreted proteins than other genes

By measuring physicochemical, compositional, and functional features for orphan and non-orphan *Meloidogyne* genes and their corresponding protein sequences, we identified features that differ between these two groups (Supplementary Table 1). Given the large size of the data set, nearly all measured features showed statistically significant differences. To identify differences that are likely to be the most biologically meaningful, we filtered the results based on relative and absolute mean differences between orphans and non-orphans.

Orphan protein sequences were generally shorter, exhibited differences in amino acid composition, especially in lysine and arginine with orphan proteins having more lysine and less arginine in their proteins, and were more positively charged at pH 7.5 compared to non-orphans. Additionally, orphan proteins were more frequently predicted to be extracellular or localized in plastids. This is consistent with the results of a previous study showing that a high lysine to arginine ratio is associated with extracellular localization ^73^.

Orphan proteins contained fewer annotated domains, as expected given their lack of detectable homologs. A notable difference concerned signal peptide prediction: 4.1% of orphan proteins were predicted to contain a signal peptide, compared to only 2.4% of non-orphan proteins. In addition, orphan proteins accounted for almost 30% of proteins with a predicted signal peptide, indicating an enrichment of signal peptides among orphan proteins. This is consistent with our observation that orphan proteins are more frequently predicted to be extracellular and correlates with our finding that nearly half of known effectors secreted by root-knot nematodes are encoded by orphan genes.

We next assessed whether orphan genes accumulate more mutations, and particularly non-synonymous ones, compared to other genes. To do so, we measured Ka and Ks values for pairs of homologous genes in pairwise comparisons of Clade III vs. Clade II *Meloidogyne* and Clade II vs. Clade I. This comparison was performed for both orphans and non-orphan genes. For orphan genes, mean values were Ka = 0.39, Ks = 2.79 and Ka/Ks = 0.19 for the comparison between Clade III and Clade II, and Ka = 0.32, Ks = 1.62 and Ka/Ks = 0.31 for the comparison between Clade II and Clade I. In contrast, for non-orphan genes, mean values were Ka = 0.12, Ks = 1.92, Ka/Ks = 0.07 and Ka = 0.21, Ks = 1.52, Ka/Ks = 0.12 for the respective comparisons. For each situation, Ka, Ks and Ka/Ks values were thus higher for orphan genes than for non-orphan genes. These results indicate that orphan genes accumulate more mutations, with a higher proportion of non-synonymous ones, suggesting accelerated evolution likely due to relaxed selection on these relatively recent genes.

### Random Forest classifiers discriminate orphan vs. non-orphan and *de novo* vs. diverged orphan proteins

Given the computational cost of our full comparative pipeline, and the observation that orphans display distinct molecular characteristics, we evaluated whether these features could be used to rapidly classify orphan genes using machine learning. We trained two Random Forest classifiers: one to distinguish orphans from non-orphans, and another to distinguish highly diverged from *de novo* orphan genes.

The orphan vs. non-orphan classifier achieved an F1 score of 0.91, with a precision of 0.94 and a recall of 0.88, indicating high overall performance and a slightly better performance to minimize the false positives. The *de novo* vs. highly diverged classifier also performed well, with an F1 score of 0.93, a precision of 0.93 and a recall of 0.92, suggesting a well-balanced and robust model for separating these two categories of orphans.

Gini impurity and shap analyses revealed that the most influential variables for classifying orphans vs. non-orphans were sequence length, probability of extracellular localization, and lysine frequency. For the *de novo* vs. highly diverged classifier, the most impactful features were mitochondrion localization probability, frequency of acidic amino acids, sequence length, and phenylalanine frequency. Although these features contributed most strongly, a wide range of additional features also influenced classification in both models, underscoring the multifactorial nature of orphans’ characteristics (Figure 7).

**Figure 7:**
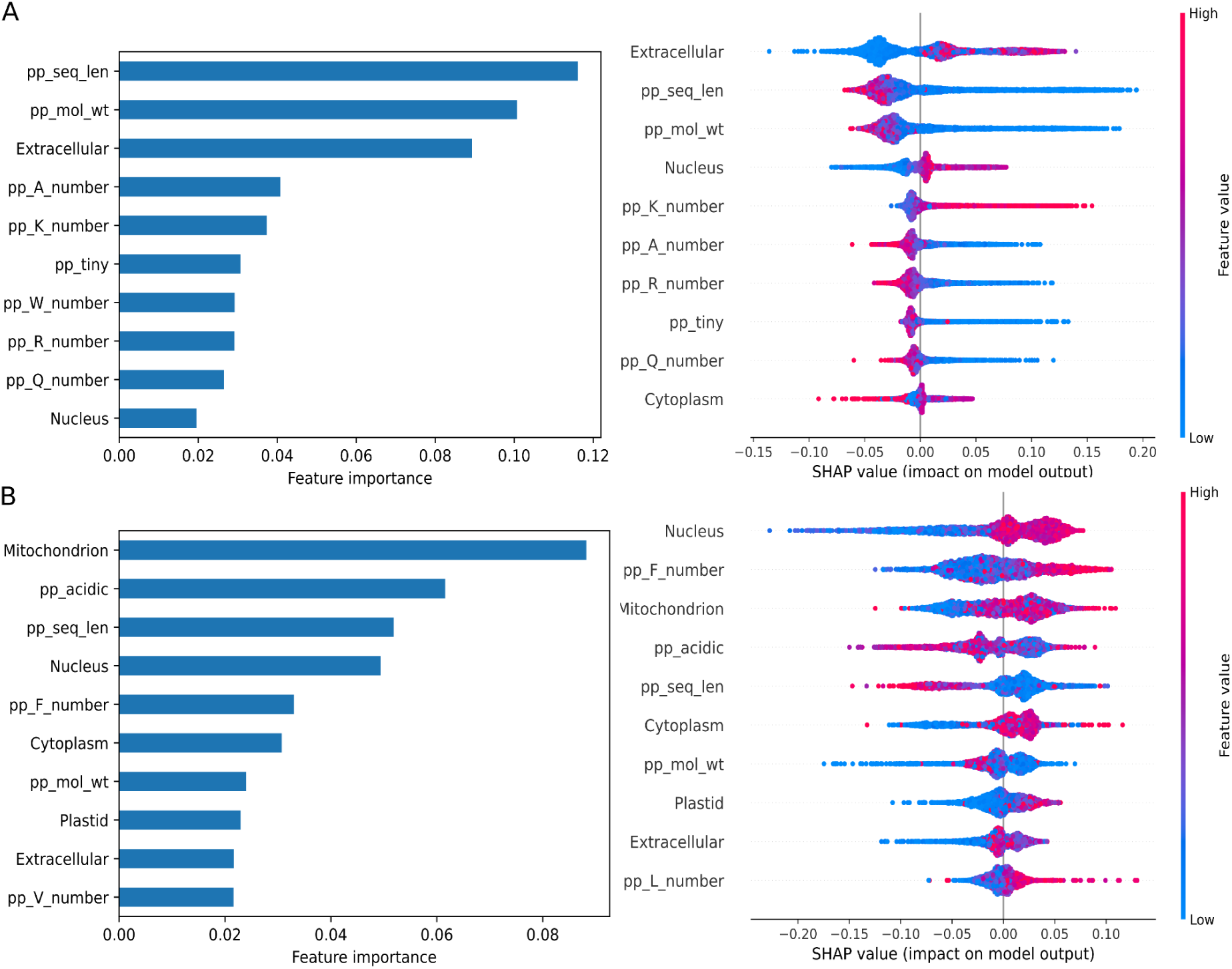
Feature importances for orphan classifier and *de novo* classifier according to Gini impurity and shap tree explainer. A. Feature importances for orphan vs. non-orphan classifier. B. Feature importances for *de novo* vs. highly diverged classifier. In each case, only the first 10 most impactful features are represented for clarity. See Supplementary Figure 5 for the plots with all the features.

We next applied the highly diverged vs. *de novo* classifier to orphans whose origin could not be determined with our pipeline. This analysis was performed for each clade. In Clade I, 56% out of the 14,807 previously unclassified orphans were predicted to be *de novo*, whereas in Clade II, only 10% out of the 4,617 unclassified orphans were predicted to be *de novo*. In Clade III, as well as among orphans universally conserved across *Meloidogyne*, only 8% out of the 6,082 unclassified orphans were predicted to be *de novo*. These results suggest a higher prevalence of *de novo* genes in Clade 1, consistent with our manual classification (Figure 5). In addition, this analysis suggests that two additional known effectors are probably encoded by *de novo* genes.

## Discussion

Since 2008, orphan genes in root-knot nematodes have attracted the attention of researchers. This interest is due to two main reasons: (i) because they are specific to *Meloidogyne* species, these genes are possibly involved in plant parasitism, and (ii), because these genes lack homologs in other species, specifically targeting them is unlikely to cause undesired side effects in non-target organisms ^32,74^. However, despite this strong interest, their origin, evolutionary history, and characteristics had remained mysterious and unexplored so far.

In this study, we provide the first comprehensive characterization of orphan genes in *Meloidogyne*, revealing their prevalence, evolutionary origins, and molecular signatures, as well as their contribution to parasitism in light of recently available genome data and advances in comparative genomics tools. We found that approximately 16% of genes in *Meloidogyne* species are transcriptionally supported, genus-specific orphan genes. This proportion is consistent with the 5-30% range previously reported for eukaryotic genomes ^1^. Direct comparison of these percentages between different studies done at different times with different methodologies remains challenging. However, the substantially high fraction we observed suggests that orphan genes are evolutionarily significant and may play important functional roles in root-knot nematodes. As a conservative strategy to avoid over-prediction by gene annotation software, we excluded species-specific orphan genes and retained only those present in at least two species. Therefore, the total number of orphan genes in root-knot nematodes is certainly higher. Furthermore, it would be interesting in future studies to confirm that some of these genes are actually species-specific, as they might represent valuable candidates for the development of markers to differentiate *Meloidogyne* species. Differentiating these species based on molecular and morphological markers remains challenging, yet is important for the farmers to adapt pest management strategies.

Our comparative analyses were performed using 85 quality-checked nematode proteomes, representing a comprehensive and diversified dataset. However, it should be kept in mind that the number of orphan genes identified remains sensitive to the completeness and diversity of available genomes. As new high-quality genomes become available, especially from lineages closely related to *Meloidogyne*, our ability to identify orphan genes and infer their origin will improve. The genus *Pratylechus* is the closest relative of *Meloidogyne* ^24–26^, and by the time we performed this comparative analysis, *Pratylenchus penetrans* was the only species with an available annotated genome meeting our completeness threshold ^71^. Although genome data have been published for three other *Pratylenchus* species, *P. coffeae* ^75^, *P. scriberni* ^76^ and *P. vulnus* ^77^, the annotation data are not publicly available for the first two and therefore could not be included. Nevertheless, including additional *Pratylenchus* species may help identify closer relatives to *Meloidogyne* and refine our inferences regarding gene novelty. This limitation is particularly relevant for our synteny-based approach, which is currently more effective within the genus *Meloidogyne*. Indeed, synteny analyses within *Meloidogyne* species, to study *de novo* gene birth within the genus, yielded more confident alignments and synteny results. In contrast, when investigating the origin of orphan genes present in the last common ancestor of *Meloidogyne* species, it was more challenging to establish a synteny around orphan genes of *Meloidogyne* with *P. penetrans* genome. This is likely due to the greater evolutionary distance between *Pratylenchus* and *Meloidogyne* compared to that between the three clades within the *Meloidogyne*. Indeed, the last common ancestor of *Meloidogyne* Clades I, II and III is estimated to be ca. 65 My old, while the last common ancestor of *Meloidogyne* and *P. penetrans* is estimated at ca. 136 My ago ^35^. This is also reflected in the lower sequence identity observed between orthologous proteins from *Pratylenchus* and *Meloidogyne,* compared with identities within *Meloidogyne*. Overall, these findings highlight the need for a better genomic representation of intermediate lineages to resolve the deeper evolutionary origins of these genes.

Using mass spectrometry and translatomic data, we were also able to verify the translation of more than half of the transcriptionally-supported orphan genes, providing further evidence that these genes are not only transcribed but also translated into proteins. However, the limited availability of proteomics data, restricted to only one *Meloidogyne* species, limits our ability to broadly validate the existence of these proteins across the genus. Indeed, mass spectrometry data were available only for *M. incognita*, meaning that translational evidence is lacking for the other Clade I species for orphans as well as non-orphans despite their large gene sets. Therefore, the proportion of translated orphan proteins reported here is likely an underestimate. Nevertheless, this analysis confirms that a substantial proportion of orphan proteins are indeed translated. Expanding proteomic coverage across *Meloidogyne* species would greatly enhance future studies.

In terms of gene origin, we estimated that high divergence from homologous genes and de novo gene birth account for 20% and 18% of orphan genes, respectively. Nevertheless, the mechanism of origin remains ambiguous for roughly half of the orphan genes, for two main reasons: (i) orphan sequences cannot always be successfully aligned to the outgroup genome, and (ii) synteny is not always conserved around the orphan gene. Moreover, gene emergence cannot always be attributable to a single mechanism, since both high divergence and de novo processes may jointly contribute to the presence of an orphan gene ^15^. However, when applying our Random Forest classifier to these ambiguous cases, we observed that *de novo* gene birth appears more prevalent in Clade I, whereas high divergence from ancestral genes is more prevalent in the other clades. This might reflect the more recent evolutionary history of Clade I species, or the importance of whole-genome duplication in these species, which could favor *de novo* gene birth. Polyploidy in these species is due to complex hybridization events ^47,48,78^, which can be associated with the re-activation and multiplication of transposable elements. Recent findings have confirmed that transposons can favor the emergence of *de novo* transcripts and, eventually, *de novo* gene birth ^72^. Another possibility is that, as most of the *de novo* genes identified by synteny analysis are in Clade I, the classifier was trained to identify them, and therefore it may be limited to identify *de novo* genes in other clades. Nonetheless, these classifiers are extremely fast compared to our full comparative genomics pipeline and unlock the possibility to scan massive datasets of predicted proteomes, offering a practical tool for rapid orphan gene identification and origin inference in future studies.

Our results also show that orphan genes, as a whole, encode proteins with distinctive features compared to non-orphan genes. Specifically, orphan proteins tend to be shorter, more positively charged, enriched in signal peptides, and have increased lysine/arginine (K/R) content, which likely explains why they tend to be more extracellular ^73^. While similar comparisons in other species identified differences in sequence length and GC content (the latter not observed here), our results highlight additional features specific to orphan proteins in root-knot nematodes ^79^. These characteristics are consistent with proteins that function in the extracellular environment and support the observation that many effectors secreted by plant-parasitic nematodes are encoded by orphan genes. The enrichment of signal peptides among orphan proteins also suggests that additional effectors remain to be discovered within the *Meloidogyne*-specific orphan gene dataset. While this hypothesis is compelling, further experimental validation is needed to determine the extent to which orphan genes contribute to the set of effector proteins and, more generally, to parasitism in these nematodes.

Overall, our study provides the first comprehensive analysis of orphan genes, including the investigation of their origin (high divergence vs. *de novo* gene emergence) and their characteristic features in *Meloidogyne*, a genus of plant-parasitic nematodes with high worldwide economic importance. We show that orphan genes constitute an important fraction of *Meloidogyne* gene sets, with a substantial proportion likely to have emerged *de novo*. These findings advance our understanding of how new genes arise and evolve in parasitic lineages and highlight the importance of orphan genes in the adaptive biology of *Meloidogyne*. Future studies should focus on the functional validation of candidate effectors among orphan genes, and on the expansion of phylogenomic and proteomic datasets to refine evolutionary inferences.

## Code Availability

The scripts written to treat the outputs of different tools as well as to treat and filter different results can be found on GitHub (repository: erseckin/melo-evo-orphan).

## Data Availability

All datasets generated in this study are publicly available in the « Orphan and *De Novo* Genes in Root-Knot Nematodes » dataverse on recherche.data.gouv.fr and have been assigned persistent DOIs.

## Acknowledgements

This research was supported by the joint INRAE-Inria PhD program, which funds the PhD thesis of Er.S. We are grateful to the genotoul bioinformatics platform Toulouse Occitanie (Bioinfo Genotoul, DOI:10.15454/1.5572369328961167E12) as well as to the bioinformatics and genomics platform, BIG, Sophia Antipolis (ISC PlantBIOs, DOI: 10.15454/qyey-ar89) for computing and storage resources. We would also like to thank Gianni Liti and Mathilde Carpentier for their suggestions and ideas that were discussed during Er.S.’s Ph.D. follow-up committee meetings.

