## Supplementary for "Orphan genes shape the genome and parasitic arsenal of root-knot nematodes"

**Table 1:** Mean average and differences of all characteristics analyzed to compare orphans and non-orphans.

A.

| Feature | Mean orphans | Mean non-orphans | Absolute mean diff | Relative mean diff | P-value |
|---|---|---|---|---|---|
| pp_seq_len | 154.3081 | 411.0554 | 256.7473 | 2.6639 | 0.0000 |
| pp_mol_wt | 17759.5517 | 46746.5820 | 28987.0304 | 2.6322 | 0.0000 |
| pp_molar_ext_coef | 16007.3062 | 42983.8628 | 26976.5566 | 2.6853 | 0.0000 |
| pp_A_number | 0.0375 | 0.0525 | 0.0149 | 1.3981 | 0 |
| pp_C_number | 0.0232 | 0.0215 | 0.0017 | 1.0808 | ~0 |
| pp_D_number | 0.0373 | 0.0467 | 0.0094 | 1.2519 | 0.0000 |
| pp_E_number | 0.0628 | 0.0681 | 0.0053 | 1.0837 | 0.0000 |
| pp_F_number | 0.0726 | 0.0526 | 0.0199 | 1.3790 | 0.0000 |
| pp_G_number | 0.0501 | 0.0548 | 0.0048 | 1.0957 | 0.0000 |
| pp_H_number | 0.0199 | 0.0206 | 0.0007 | 1.0373 | 0.0000 |
| pp_I_number | 0.0787 | 0.0709 | 0.0078 | 1.1095 | 0.0000 |
| pp_K_number | 0.0861 | 0.0703 | 0.0158 | 1.2243 | 0.0000 |
| pp_L_number | 0.0975 | 0.0956 | 0.0018 | 1.0192 | ~0 |
| pp_M_number | 0.0259 | 0.0225 | 0.0034 | 1.1519 | ~0 |
| pp_N_number | 0.0685 | 0.0632 | 0.0053 | 1.0838 | ~0 |
| pp_P_number | 0.0421 | 0.0444 | 0.0023 | 1.0547 | 0.0000 |
| pp_Q_number | 0.0357 | 0.0448 | 0.0092 | 1.2573 | 0.0000 |
| pp_R_number | 0.0443 | 0.0517 | 0.0073 | 1.1654 | 0.0000 |
| pp_S_number | 0.0769 | 0.0759 | 0.0009 | 1.0123 | ~0 |
| pp_T_number | 0.0474 | 0.0503 | 0.0029 | 1.0609 | 0.0000 |
| pp_V_number | 0.0475 | 0.0509 | 0.0033 | 1.0701 | 0.0000 |
| pp_W_number | 0.0101 | 0.0115 | 0.0014 | 1.1353 | 0.0000 |
| pp_Y_number | 0.0360 | 0.0312 | 0.0048 | 1.1547 | ~0 |
| pp_acidic | 0.1001 | 0.1148 | 0.0147 | 1.1464 | 0.0000 |
| pp_aliphatic | 0.2612 | 0.2699 | 0.0087 | 1.0332 | ~0 |

| | | | | | |
|---|---|---|---|---|---|
| pp_aromatic | 0.1386 | 0.1159 | 0.0227 | 1.1954 | 0.0000 |
| pp_basic | 0.1503 | 0.1426 | 0.0077 | 1.0540 | ~0 |
| pp_charge_at_pH | 1.6785 | -0.1075 | 1.7860 | -15.6102 | ~0 |
| pp_charged | 0.2505 | 0.2574 | 0.0070 | 1.0277 | ~0 |
| pp_gravy | -0.2500 | -0.3583 | 0.1082 | 0.6978 | ~0 |
| pp_helix | 0.3424 | 0.3127 | 0.0297 | 1.0949 | 0.0000 |
| pp_instab_idx | 43.7761 | 45.4205 | 1.6444 | 1.0376 | ~0 |
| pp_isoelec_point | 8.0979 | 7.5389 | 0.5590 | 1.0741 | 0.0000 |
| pp_mean_kd_hydro | -0.2681 | -0.3653 | 0.0972 | 0.7338 | ~0 |
| pp_mean_rose_hydro | 0.7255 | 0.7220 | 0.0035 | 1.0049 | ~0 |
| pp_mean_vihinen_flex | 0.9997 | 1.0016 | 0.0019 | 1.0019 | ~0 |
| pp_non_polar | 0.5211 | 0.5083 | 0.0128 | 1.0251 | ~0 |
| pp_polar | 0.4789 | 0.4917 | 0.0128 | 1.0267 | ~0 |
| pp_sheet | 0.2237 | 0.2386 | 0.0149 | 1.0668 | 0.0000 |
| pp_small | 0.5211 | 0.5083 | 0.0128 | 1.0251 | ~0 |
| pp_tiny | 0.2351 | 0.2550 | 0.0199 | 1.0848 | 0.0000 |
| pp_turn | 0.2375 | 0.2383 | 0.0009 | 1.0036 | ~0 |
| A_frequency | 0.3423 | 0.3388 | 0.0035 | 1.0103 | ~0 |
| C_frequency | 0.1582 | 0.1580 | 0.0001 | 1.0009 | ~0 |
| G_frequency | 0.1801 | 0.1883 | 0.0082 | 1.0454 | 0 |
| T_frequency | 0.3194 | 0.3149 | 0.0045 | 1.0144 | ~0 |
| AT_frequency | 0.6617 | 0.6537 | 0.0080 | 1.0123 | ~0 |
| GC_frequency | 0.3383 | 0.3463 | 0.0080 | 1.0237 | ~0 |
| Cell membrane | 0.1796 | 0.2279 | 0.0483 | 1.2692 | ~0 |
| Endoplasmic reticulum | 0.2929 | 0.2325 | 0.0604 | 1.2597 | 0.0000 |
| Extracellular | 0.3723 | 0.1821 | 0.1901 | 2.0439 | 0.0000 |
| Golgi apparatus | 0.2193 | 0.2120 | 0.0073 | 1.0344 | 0.0000 |
| Lysosome/Vacuole | 0.2442 | 0.1987 | 0.0455 | 1.2291 | 0.0000 |
| Mitochondrion | 0.2529 | 0.2114 | 0.0414 | 1.1959 | 0.0000 |

| | | | | | |
|---|---|---|---|---|---|
| Nucleus | 0.4108 | 0.3975 | 0.0133 | 1.0334 | ~0 |
| Peroxisome | 0.0436 | 0.0376 | 0.0060 | 1.1600 | 0.0000 |
| Plastid | 0.0415 | 0.0303 | 0.0112 | 1.3707 | 0.0000 |

B.

| Feature | P-value | Contigency table |
|---|---|---|
| InterPro domain | 0 | {0: {0: 41585, 1: 45676}, 1: {0: 161720, 1: 3927}} |
| Peptide signal | ~0 | {0: {0: 198415, 1: 47542}, 0: {0: 4890, 1: 2061}} |

A. Means and significance of mean differences B. Contingency table for categorical features where 0 means absence and 1 means presence for each state. Protein characteristics: protein sequence length, molecular weight, IInstability index (calculated according to Guruprasad et al, 1990), Average flexibility value (calculated according to Vihinen, 1994), Average hydrophobicity profile value (calculated according to Kyte and Doolittle, 1982), The average value of hydropathy (calculated according to Kyte and Doolittle, 1982), The isoelectric point and charge of the protein at pH 7.5 (calculated by Bio.SeqUtils.ProtParam), The calculation of the molar extinction coefficient with respect to cysteine residues and cysteine-cysteine bonds (calculated by Bio.SeqUtils.ProtParam). The % of small amino acids (corresponds to the cumulative percentage of amino acids A, C, G, S, T),  The % of aliphatic amino acids (corresponds to the cumulative percentage of amino acids A, I, L, V), % aromatic amino acids (Cumulative percentage of amino acids F, H, W, Y), % of non-polar amino acids (Cumulative percentage of amino acids A, C, F, G, I, L, M, P, V, W, Y),  % of polar amino acids ( Cumulative percentage of D, E, H, K, N, Q, R, S, T, Z amino acids), % of charged amino acids: (Cumulative percentage of amino acids B, D, E, H, K, R, Z), % basic amino acids (Cumulative percentage of amino acids H, K , R), % of acidic amino acids (Cumulative percentage of B, D, E, Z amino acids), % of amino acids in a helix, Cumulative percentage of amino acids that tend to be found in a helix: F, I, Y, V, W, L), % of amino acids in a loop, Cumulative percentage of amino acids that tend to occur in a loop: N, P, G, S), % of amino acids in a leaflet, cumulative percentage of amino acids that tend to occur in a leaflet: E, M, A, L), % amino acid A, % amino acid C, % amino acid D, % amino acid E, % amino acid F, % amino acid G, % amino acid H, % amino acid I, % amino acid K, % amino acid L, % amino acid M, % amino acid N, % amino acid P, % amino acid Q, % amino acid R, % amino acid S, % amino acid T, % amino acid V, % amino acid W, % amino acid Y,  domain frequency (calculated by InterProScan), fPeptide signal frequency, The frequency of presence of a signal peptide for the protein sequence), as well as subcellular localization calculated by DeepLoc (probability of being in the nucleus, probability of being in the cytoplasm, probability of being extracellular, probability of being in the membrane, probability of being in the mitochondrion, probability of being in the plastid, probability of being in the ER, probability of being in the lysosome, probability of being in the golgi, probability of being the peroxisome). Gene characteristics: % of A nucleotide, % of T nucleotide, % of G nucleotide, % of C nucleotide, % of AT and % GC cumulative.

**Table 2 :** .xlsx file with available nematode genomes (species_table.xlsx).

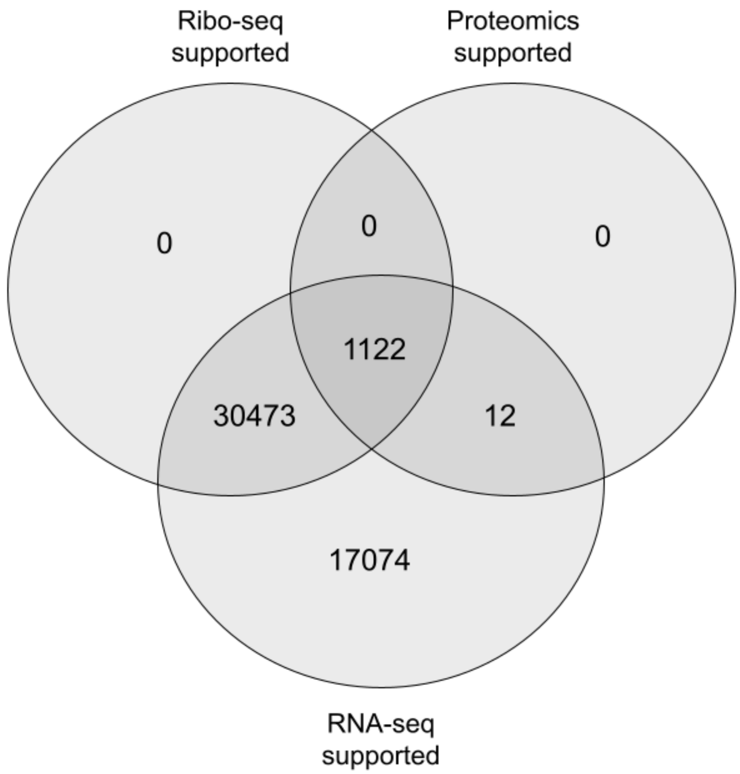


**Figure 1:** Venn diagram summarizing transcriptional and translational evidences for identified orphan proteins

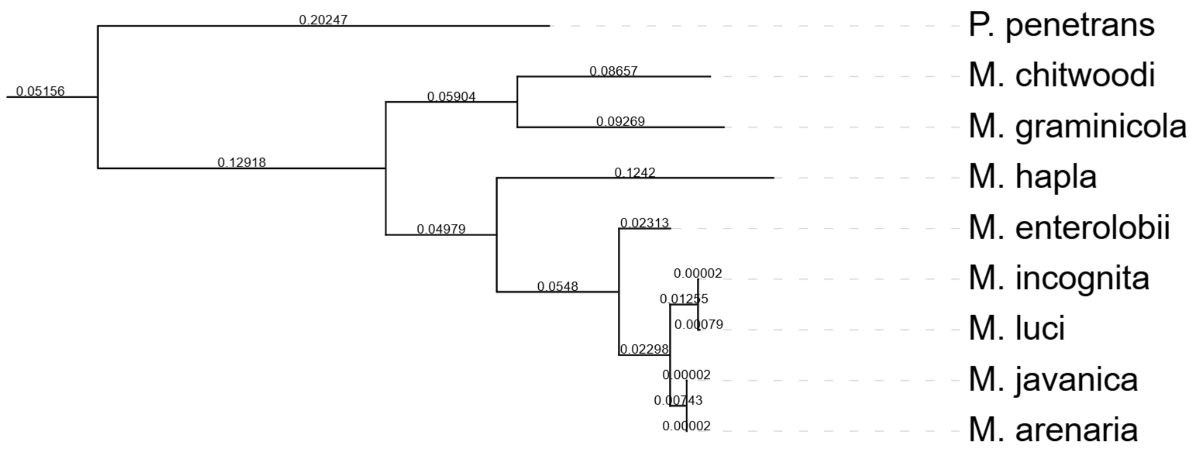


**Figure 2:** Inferred species tree by Orthofinder for *Meloidogyne* genus and its closest outgroup *P. penetrans*

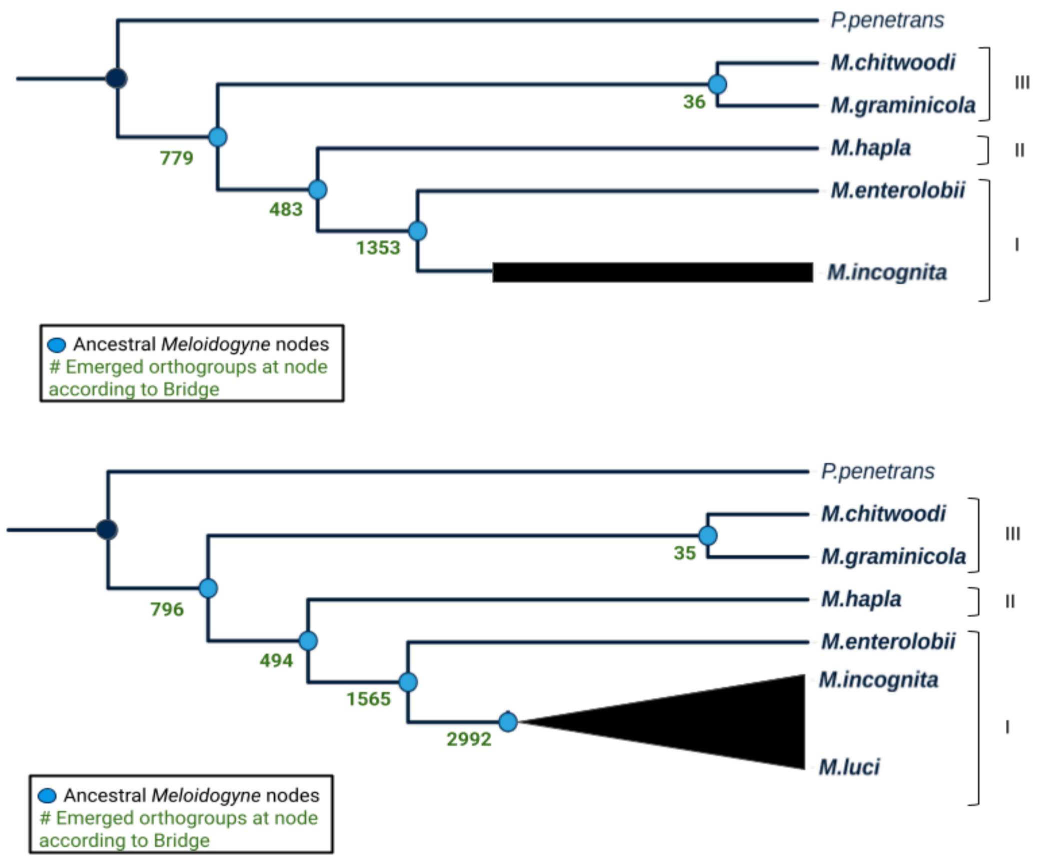


**Figure 3**: Number of emerged orphan orthogroups when species are subtracted from Clade I

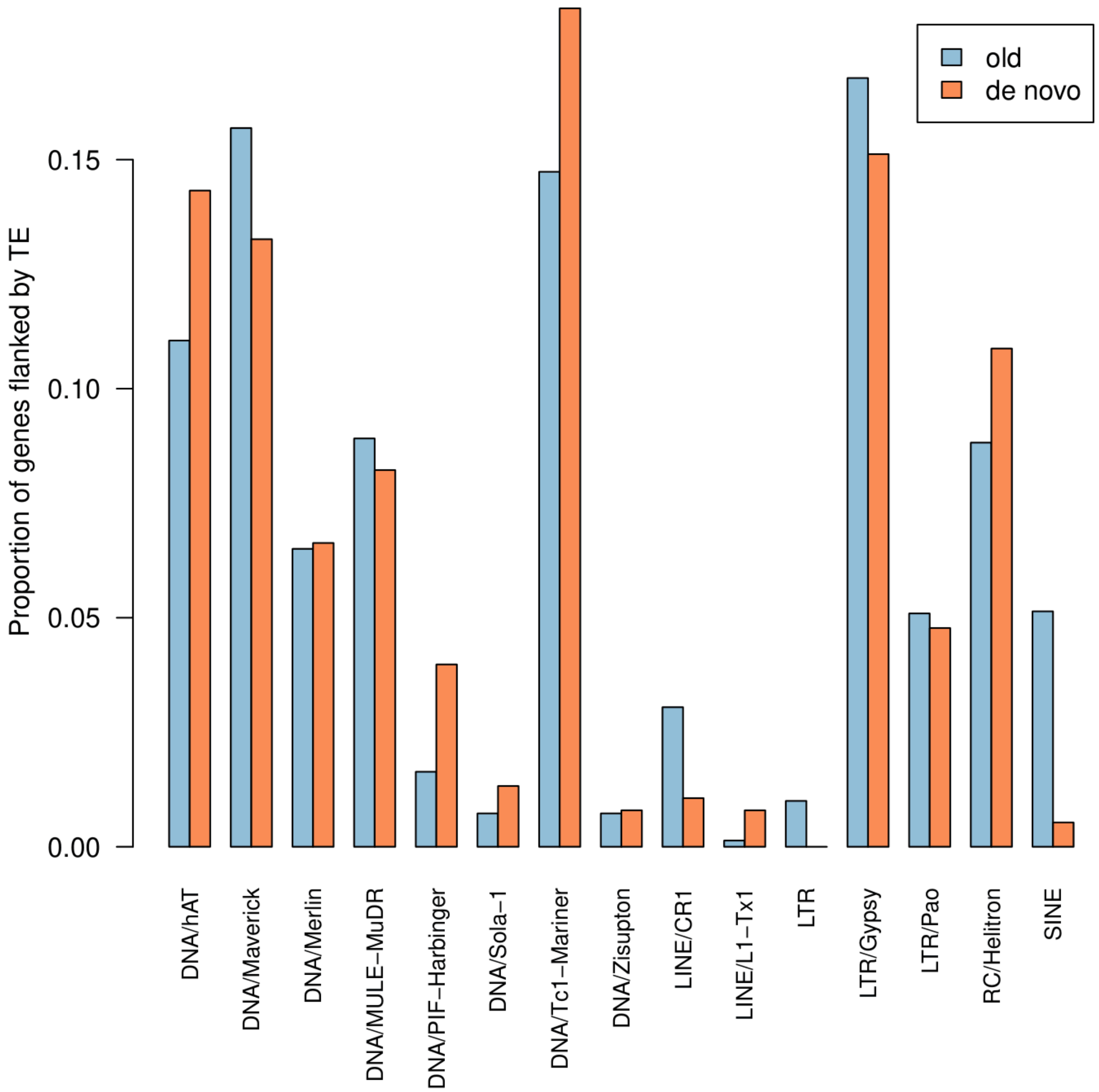


**Figure 4:** Proportion of *de novo* genes and old genes (commun to nematoda and tartigrades) flanked by transposable elements.

A.

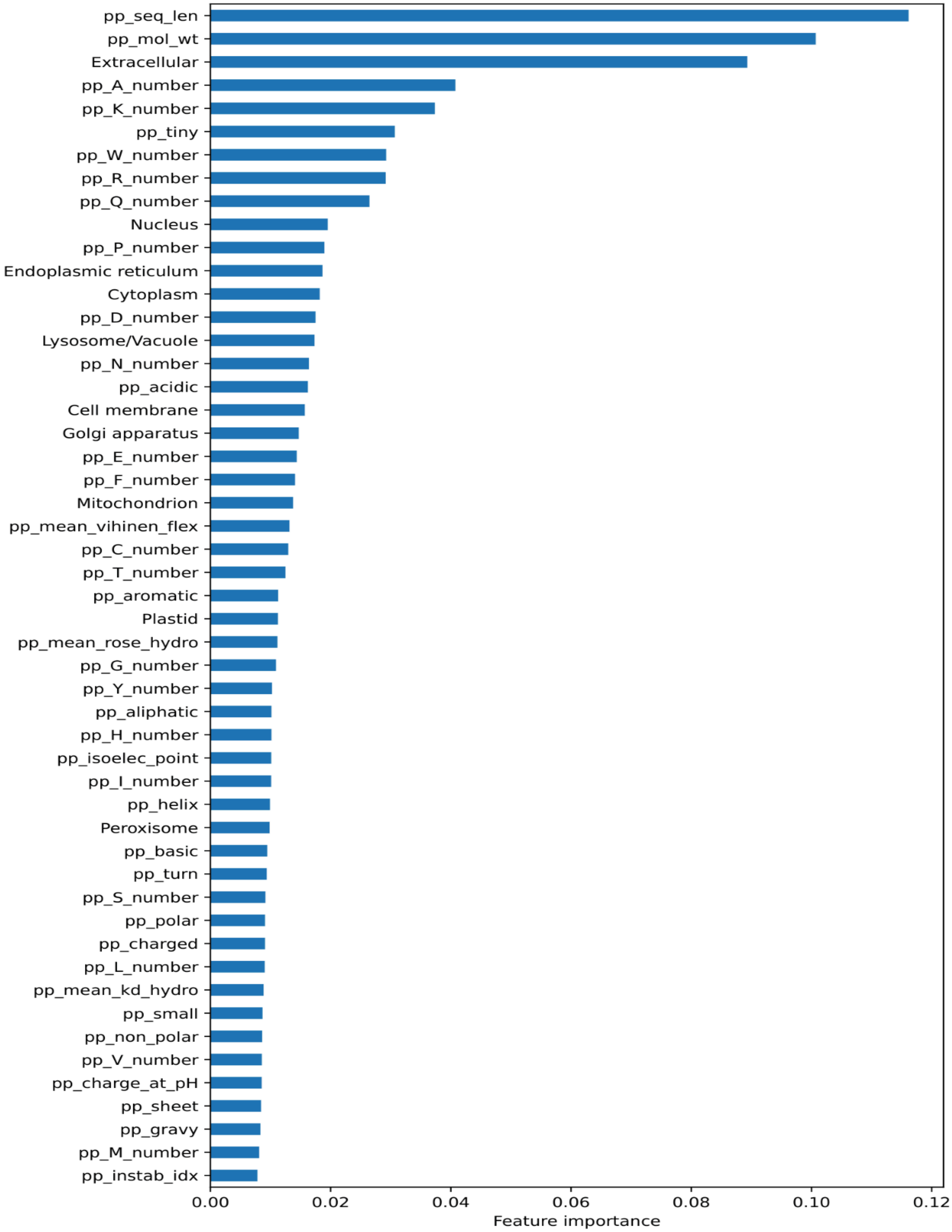

pp_seq_len
pp_mol_wt
Extracellular
pp_A_number
pp_K_number
pp_tiny
pp_W_number
pp_R_number
pp_Q_number
Nucleus
pp_P_number
Endoplasmic reticulum
Cytoplasm
pp_D_number
Lysosome/Vacuole
pp_N_number
pp_acidic
Cell membrane
Golgi apparatus
pp_E_number
pp_F_number
Mitochondrion
pp_mean_vihinen_flex
pp_C_number
pp_T_number
pp_aromatic
Plastid
pp_mean_rose_hydro
pp_G_number
pp_Y_number
pp_aliphatic
pp_H_number
pp_isoelec_point
pp_I_number
pp_helix
Peroxisome
pp_basic
pp_turn
pp_S_number
pp_polar
pp_charged
pp_L_number
pp_mean_kd_hydro
pp_small
pp_non_polar
pp_V_number
pp_charge_at_pH
pp_sheet
pp_gravy
pp_M_number
pp_instab_idx
0.00
0.02
0.04
0.06
0.08
0.10
0.12
Feature importance

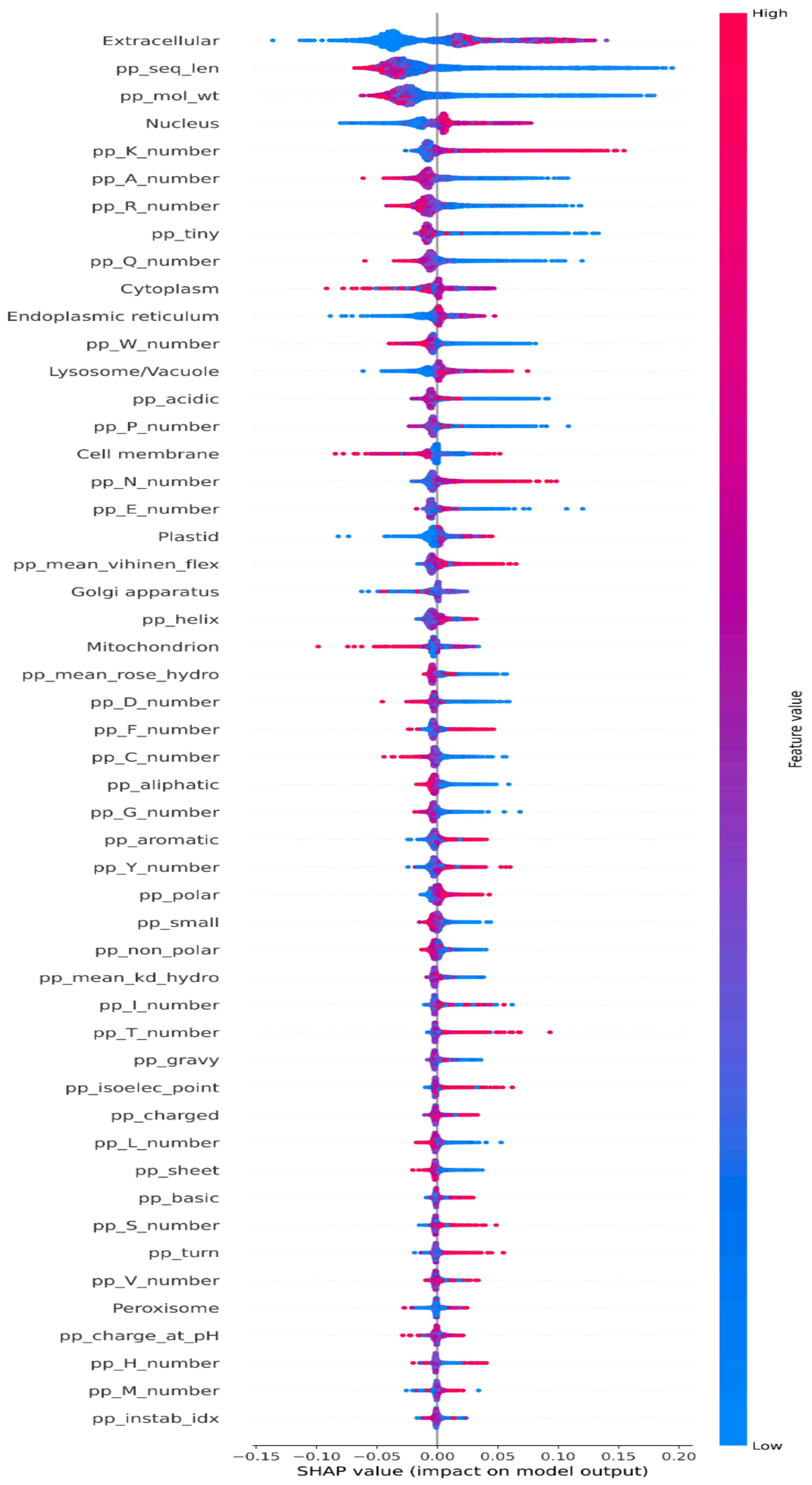

Extracellular
pp_seq_len
pp_mol_wt
Nucleus
pp_K_number
pp_A_number
pp_R_number
pp_tiny
pp_Q_number
Cytoplasm
Endoplasmic reticulum
pp_W_number
Lysosome/Vacuole
pp_acidic
pp_P_number
Cell membrane
pp_N_number
pp_E_number
Plastid
pp_mean_vihinen_flex
Golgi apparatus
pp_helix
Mitochondrion
pp_mean_rose_hydro
pp_D_number
pp_F_number
pp_C_number
pp_aliphatic
pp_G_number
pp_aromatic
pp_Y_number
pp_polar
pp_small
pp_non_polar
pp_mean_kd_hydro
pp_I_number
pp_T_number
pp_gravy
pp_isoelec_point
pp_charged
pp_L_number
pp_sheet
pp_basic
pp_S_number
pp_turn
pp_V_number
Peroxisome
pp_charge_at_pH
pp_H_number
pp_M_number
pp_instab_idx
−0.15 −0.10 −0.05 0.00 0.05 0.10 0.15 0.20
SHAP value (impact on model output)
High
Low
Feature value

B.

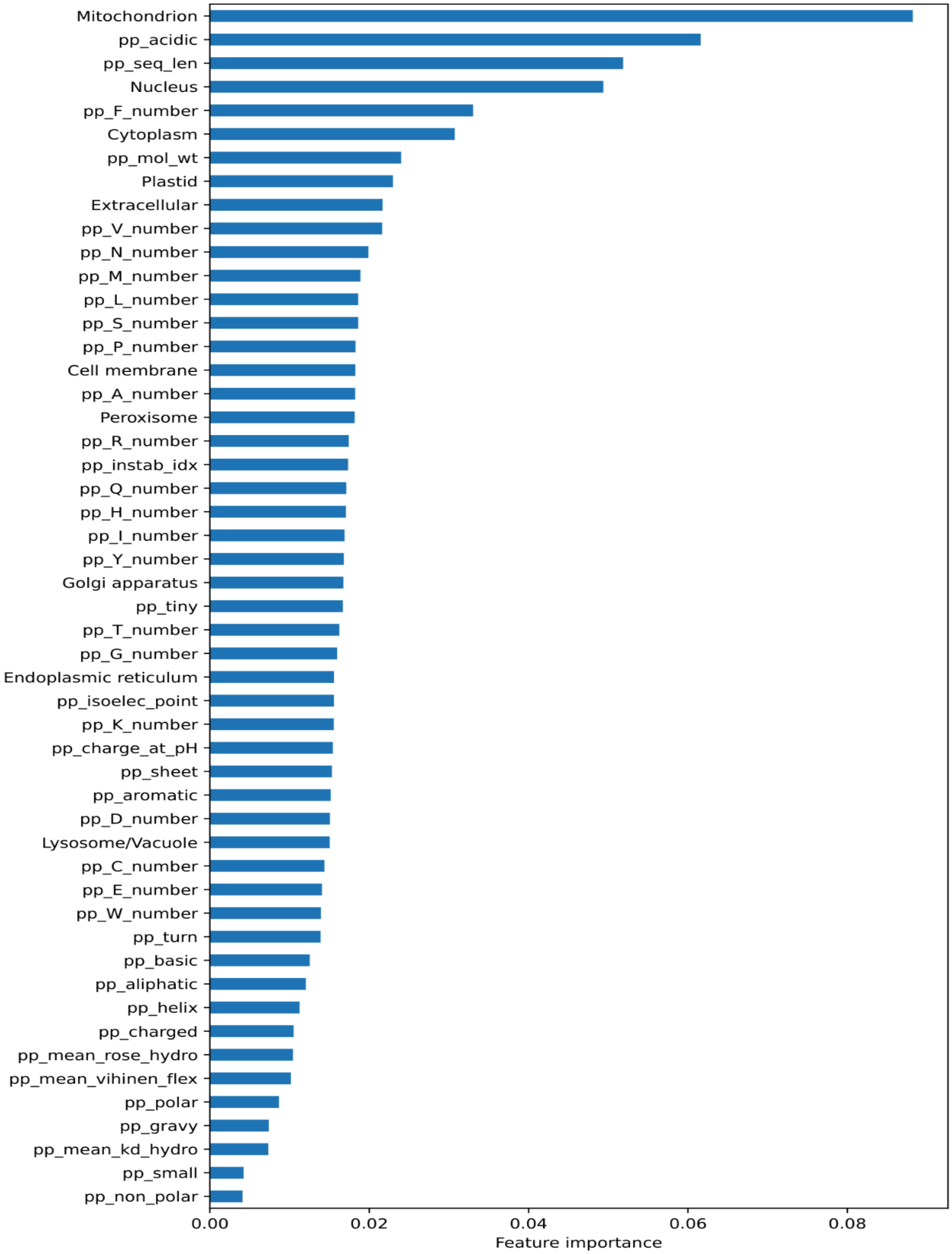
Mitochondrion
pp_acidic
pp_seq_len
Nucleus
pp_F_number
Cytoplasm
pp_mol_wt
Plastid
Extracellular
pp_V_number
pp_N_number
pp_M_number
pp_L_number
pp_S_number
pp_P_number
Cell membrane
pp_A_number
Peroxisome
pp_R_number
pp_instab_idx
pp_Q_number
pp_H_number
pp_I_number
pp_Y_number
Golgi apparatus
pp_tiny
pp_T_number
pp_G_number
Endoplasmic reticulum
pp_isoelec_point
pp_K_number
pp_charge_at_pH
pp_sheet
pp_aromatic
pp_D_number
Lysosome/Vacuole
pp_C_number
pp_E_number
pp_W_number
pp_turn
pp_basic
pp_aliphatic
pp_helix
pp_charged
pp_mean_rose_hydro
pp_mean_vihinen_flex
pp_polar
pp_gravy
pp_mean_kd_hydro
pp_small
pp_non_polar
0.00
0.02
0.04
0.06
0.08
Feature importance

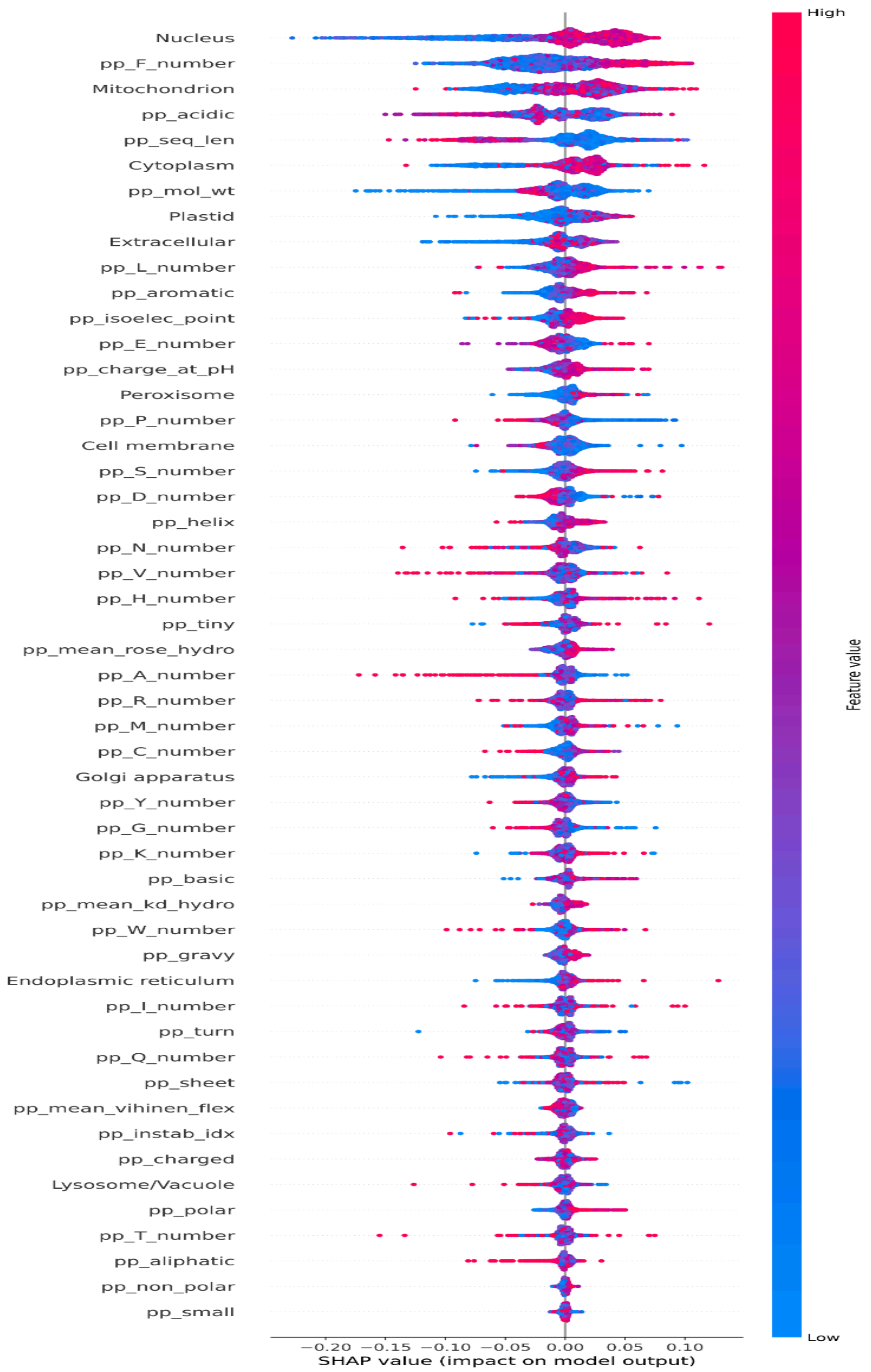


**Figure 5:** Feature importances with all features according to Gini impurity and Shap. A. For orphan vs non-orphan classifier. B. For highly diverged vs *de novo* classifier.
